# Large-scale 3D stitching montage cryo-electron tomography reveals heterogeneity and connectivity of lamellipodial actin networks

**DOI:** 10.64898/2026.11.19.767786

**Authors:** Manjunath G Javoor, Kain van den Elsen, Christoph Sommer, Robert Hauschild, Victor-Valentin Hodirnau, Noah Weber, Benedikt Singer, Michael Sixt, Florian KM Schur

**Author notes:** These authors contributed equally to this work (Christoph Sommer, Robert Hauschild).

## Abstract

Cellular processes range from local interactions among a few proteins to micrometer-scale assemblies whose coordinated components give rise to emergent properties. A prime example is the actin cytoskeleton in protrusive structures at the leading edge of migrating cells, termed lamellipodia. Significant progress has been made in understanding the functional contributions of distinct actin-filament populations and associated proteins within lamellipodia. Yet, a detailed quantitative appreciation of how actin filaments form a spatially connected and interdependent network has been impeded by limitations of existing methods. Cryo-electron tomography (cryo-ET) can visualize such biological entities at molecular resolution in native cellular environments, but its limited field-of - view constrains quantitative analysis of extended assemblies. Hence, how lamellipodial actin architectures collectively support productive migration is not completely understood. Here, we present a 3D stitching montage cryo-ET workflow that advances both methodological and biological frontiers through its application to lamellipodia. We combine large-area montage-tilt series acquisition with independent tile reconstruction, denoising, and optical flow-based three-dimensional stitching to reconstruct large regions of the leading edge up to ∼ 100 μm² at single-filament resolution. Bridging molecular resolution and cellular-scale context allows comprehensive filament tracing and quantification of actin filament networks, providing parameters for models of force generation in motile cells. Our data reveal that the network architecture in a seemingly uniform protrusive lamellipodium is markedly heterogeneous. By mapping Arp2/3 branch junction-based connectivity, using the molecular resolution of our data, we show how branch-junction distribution and network connectivity vary as a function of filament density and distance from the leading edge. We also identify the complex coexistence of short, branched and long, unbranched filament assemblies. We provide montage 3D volumes and filament vectorizations through an interactive, open web-based resource for browsing and interrogating the data. Our work thus provides a quantitative reference for lamellipodial actin architecture and a scalable approach for large-scale cryo-ET.

## Introduction

Biological processes within cells depend on the coordinated interplay of numerous components, spanning local interactions involving only a few proteins to micrometer-scale assemblies. Structural visualization of such processes has long been a key experimental step toward understanding their function. For large multicomponent assemblies, cryo-electron tomography (cryo-ET) has emerged as method of choice for resolving macromolecular structures within natively preserved cellular environments (Beck & Baumeister, 2016; Young & Villa, 2023). By combining molecular-scale structural information with cellular context, cryo-ET can reveal molecular abundance, spatial organization, and higher-order relationships that are not captured by analyses of isolated components.

A fundamental trade-off in cryo-ET, however, is the need to balance achievable spatial resolution with the desired field-of-view (FOV). This limitation becomes particularly important for biological systems whose function emerges from structural organization over length scales substantially larger than those captured by conventional high-resolution cryo-ET, which usually lie in the range of 1–3 μm².

A key example of such a biological system is the actin cytoskeleton and its role in cell migration, where motility is driven by the dynamic polymerization of globular actin (G-actin) into higher-order filamentous actin (F-actin) architectures that generate functionally distinct structures with characteristic network geometries. Lamellipodia are a model protrusion that have been extensively studied using functional and microscopy-based approaches to understand actin-network features at both the protein and architectural levels (Fäßler, Javoor, & Schur, 2023; Small et al., 2002). These 150-200nm thin, sheet-like protrusions form at the leading edge of cells and can reach sizes of more than 100 μm².

Functional studies perturbing individual actin-binding proteins have shown that, while lamellipodia ultimately depend on Arp2/3-mediated actin nucleation to generate branched networks of defined geometry (Anderson et al., 2017; Fäßler, Javoor, Datler, et al., 2023; Steffen et al., 2013; Suraneni et al., 2012; Svitkina & Borisy, 1999; C. Wu et al., 2012), the loss of other actin regulators, including formins (Kage et al., 2017; C. Yang et al., 2007), points to the existence of distinct populations of, for example, long filaments that contribute disproportionately to protrusive force generation and lamellipodial ultrastructure. Despite their morphologically simple appearance, lamellipodia therefore cannot be regarded as homogeneous sheets of branched actin. Instead, filament populations with distinct ultrastructural and dynamic properties coexist within the same protrusion and collectively shape the emergent behaviour of the lamellipodium (Blanchoin et al., 2014; Boiero Sanders et al., 2020). Specifically, fluorescence-recovery after photobleaching (FRAP) experiments of lamellipodia suggested that while most actin filaments turn over rapidly, some filaments might display increased stability and lifetime within the network (Aroush et al., 2017). In this regard, it is unclear how network parameters, including total filament number and orientation, filament length distribution, branching, connectivity, and local filament density, vary spatially within lamellipodia. Moreover, it is unclear how these geometrical properties relate to the collective behavior of the actin network, resulting in spatially adjacent and intermixing regions that protrude, retract or stall.

A cellular-scale quantitative description of actin-network architecture at single-filament resolution would be crucial to test competing or complementary biophysical models of force generation, which make different assumptions about where polymerization and depolymerization occur, how new filaments are generated, how actin architecture is spatially patterned, and how force is transmitted to the membrane (Aroush et al., 2017; J. Lee et al., 1993; Mogilner & Oster, 1996; Mueller et al., 2017; Ponti et al., 2004; Watanabe & Mitchison, 2002). The contested lamellipodium–lamella/two-layer model contains parts that are compatible with aspects of a functional multi-network hypothesis, as it predicts the coexistence of distinct filament populations that differ in their polymerization dynamics, turnover kinetics, molecular composition, and ultrastructural organization (Danuser, 2009; Giannone et al., 2007; Ponti et al., 2004; Vallotton & Small, 2009). Such filament heterogeneity could provide the lamellipodial actin network with the architectural and mechanical versatility required to support different emergent migratory properties, including variations in protrusion rate and persistence, force generation, adhesion coupling, and directional migration.

Historically, actin-based protrusions were visualized at single-filament resolution mainly by two-dimensional electron microscopy following extensive chemical preparation, limiting reliable and quantitative analysis (Koestler et al., 2008; Svitkina & Borisy, 1999). Cryo-ET later enabled 3D visualization of the actin cytoskeleton, but such datasets only provided local snapshots creating a fundamental sampling problem (Fäßler, Javoor, Datler, et al., 2023; Marston et al., 2019; Vinzenz et al., 2012). A single tomogram may contain a highly branched region, bundled region, and actively protruding or retracting edge, without revealing whether that architecture is representative of the protrusion or reflects only one of several coexisting network states. The limited FOV also truncates long filaments and disconnects filament architectures, preventing accurate measurements of filament continuity, topology, and network-scale connectivity.

Overcoming this limitation requires imaging approaches that bridge molecular resolution and cellular-scale context. Montage cryo-ET, i.e., stitching individually acquired regions into a continuous large FOV, expands the accessible area while preserving the resolution required to trace individual actin filaments. However, existing montage cryo-ET workflows generally rely on stitching two-dimensional projections before tomographic reconstruction (Brown et al., 2026; Chua et al., 2024, 2026; Hylton et al., 2026; Peck et al., 2022; J. E. Yang et al., 2023, 2026), potentially propagating errors from local sample deformation, geometric inconsistencies, or imperfect alignment into the final volume. In addition, the resulting large, stitched datasets can impede downstream processing, including recently developed denoising approaches. Specifically, the precise stitching to the accuracy of features of individual molecules, e.g. ensuring continuity of fine details such as individual filaments within dense actin networks, represents a challenge. Previous montage cryo-ET implementations have not comprehensively validated the stitching of the tiles.

Here, we have developed a montage cryo-ET workflow that combines large-area acquisition with independent tomographic reconstruction and seamless 3D stitching using optical flow-based methods, allowing also to image and reconstruct irregular sample areas of any shape (Figure 1 A1-A3). This enabled us to visualize cellular protrusions over large areas while maintaining the resolution required to trace individual actin filaments. Because reconstructed tiles remain accessible both independently and within the assembled volume in different processing states, the workflow supports local high-resolution processing while preserving the continuous spatial context required for network-scale analysis. We further combine these reconstructions with quantitative filament tracing and analysis across extended regions of the leading edge (Figure 1 A4). Importantly, this allows local filament architectures to be interpreted within the context of the entire protrusion and enables spatially restricted network states to be distinguished from properties characteristic of the lamellipodium as a whole.

**Figure 1:**
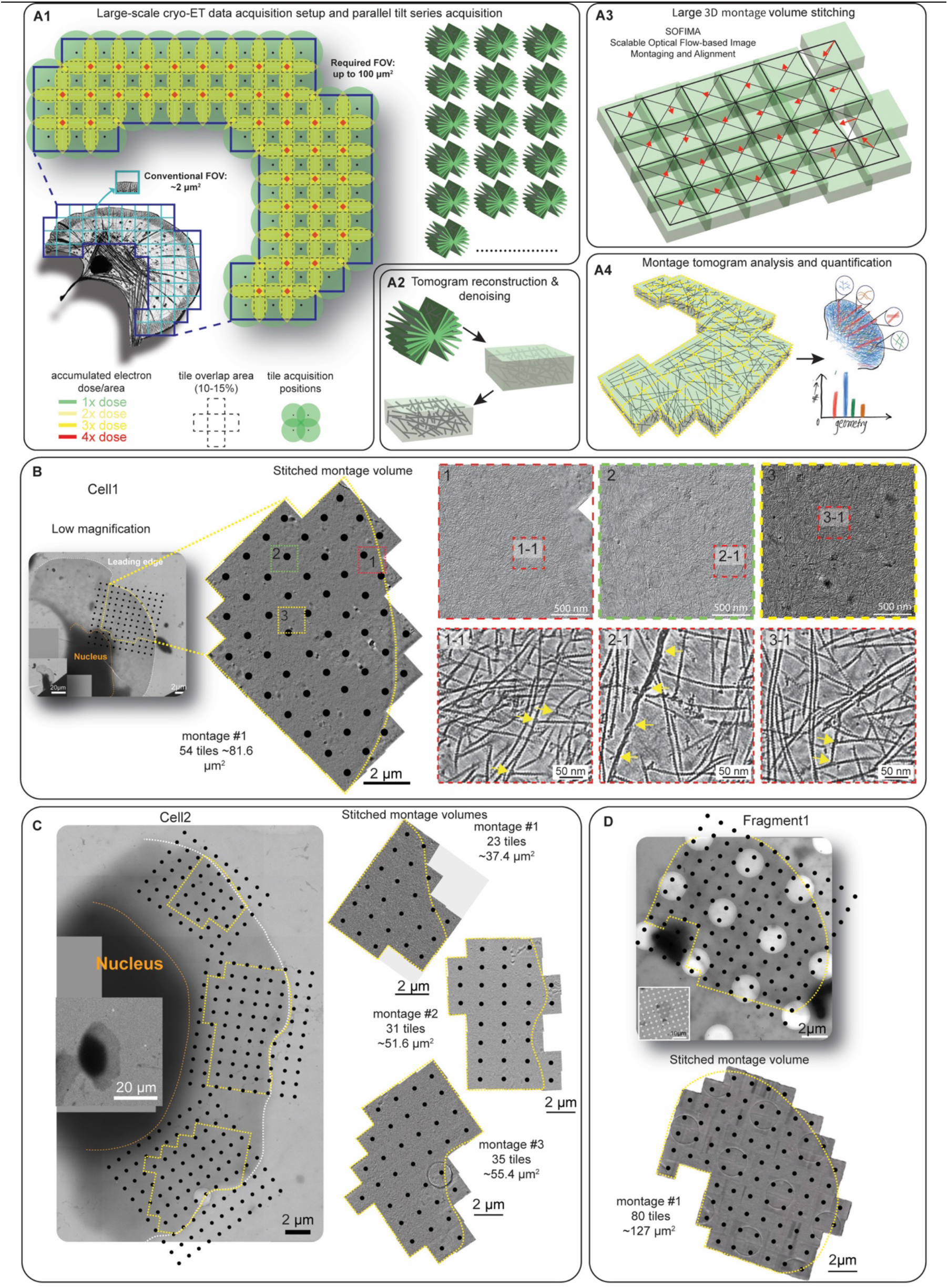
Workflow for large-area montage cryo-electron tomography of keratocyte actin networks. **A1**. Large-scale cryo-ET acquisition strategy: Regions of interest larger than the conventional single-tomogram field of view are covered by overlapping montage tiles, with parallel tilt-series acquisition at each tile position. Tile overlap, beam size, and acquisition geometry is optimized to balance stitching accuracy with accumulated electron dose. **A2**. Individual tilt series are reconstructed into tomograms and denoised to improve actin filament visibility. **A3**. Overlapping tomograms are stitched into continuous large-volume reconstructions using three-dimensional, optical flow-based alignment in SOFIMA. **A4**. The stitched montage tomograms are used for filament tracing, validation of filament continuity across tile boundaries, and quantitative analysis of actin network organization. **(B-D)** Montage acquisitions on two keratocytes and one keratocyte fragment. In the low magnification overview 2D maps, black dots mark the PACE-tomo acquisition points, yellow dotted outlines mark the acquired regions, the leading edge (white dotted line) and nucleus (orange dotted line) are annotated. Overviews are shown for all presented examples as 1.352-nm slices through the cryo-CARE denoised, stitched montage volumes. For panel B increased magnification views are shown to illustrate data quality and detail. **(B)** Cell1. From left to right: low magnification overview 2D map; stitched volume of montage#1 (54 tiles, ∼81.6 µm²), showing seamless stitching; higher-magnification views (1–3) of the red, green, and yellow boxed regions, showing filament continuity; and enlarged views (1-1, 2-1, 3-1) of the boxed regions in 1–3, showing actin filaments at high resolution. Arrows mark branch junctions, and intermediate filaments. Scale bars: 20 µm (inset); 2 µm (low magnification map and stitched volume); 500 nm (1–3); 50 nm (1-1, 2-1, 3-1). **(C)** Cell2. Left: low magnification overview 2D map showing the three regions for which montages were acquired. Right: stitched volumes of montage#1 (23 tiles, ∼37.4 µm²), montage#2 (31 tiles, ∼51.6 µm²), and montage#3 (35 tiles, ∼55.4 µm²). Scale bars: 20 µm (inset); 2 µm (overview and stitched volumes). **(D)** Keratocyte fragment#1. Top: low magnification overview 2D map. Bottom: stitched volume (80 tiles, ∼127 µm²). Scale bars: 10 µm (inset); 2 µm (selected-area map and stitched volume).

## Results

### Fish keratocyte sample preparation

Fish epidermal keratocytes are well established models to study lamellipodia-dependent cell motility. They are among the fastest and most persistently moving cells, exhibiting a broad, thin lamellipodium at their leading edge (Keren et al., 2008; J. Lee et al., 1993). Their overall homogenous cell morphology and continuous movement with a speed of tens of micrometers per minute eliminates the need for correlative imaging to distinguish strongly varying protrusive migration states, while local variations at the leading edge still allow analysis of subtle differences in protrusion dynamics and architecture. In addition, due to their ability to form - upon treatment with the kinase inhibitor staurosporine (Euteneuer & Schliwa, 1984; Verkhovsky et al., 1999) - nucleus-free cytoplasmic fragments that retain the core motility machinery, we considered them an ideal system for establishing our large-scale cryo-ET workflow to study the leading-edge actin network (Figure S1A).

Given the importance of i) proper cell spreading on EM grid substrates, ii) achieving a sufficient signal-to-noise ratio (SNR), and iii) minimizing sensitivity to electron dose damage, we optimized the seeding of keratocyte cells and fragments on grids with a silicon dioxide (SiO₂) substrate layer (Figure S1B, C), which we found to outperform amorphous carbon substrate both in terms of mechanical stability and electron transparency. To further facilitate the resolution of the dense lamellipodial actin network by cryo-ET, we employed an established protocol combining membrane extraction and fixation prior to vitrification, which has been shown to reliably preserve cytoskeletal architecture, including actin filaments and associated binding proteins, such as the Arp2/3 complex (Fäßler, Dimchev, et al., 2020; Fäßler et al., 2023). Specifically, using this protocol we obtained a 3.6 Å structure of zebrafish F-actin within the cellular actin network using single-particle cryo-EM, demonstrating the preservation of high-resolution actin filament structure (Table S1, Figure S1D, E). With these settings established, we proceeded to implement montage cryo-ET data acquisition of both cytoplasmic fragments and intact cells on holey (2 µm holes spaced 2 µm apart) and semi-continuous SiO₂ substrates (1 µm holes spaced 20 µm apart), respectively.

### Establishment of large-scale montage cryo-ET data acquisition

To meet the requirement of achieving seamless FOVs approximating the size of lamellipodia while retaining sufficient resolution to clearly resolve actin filaments (∼7–8 nm diameter) and bound proteins, we selected a pixel size approximately 25-fold smaller than the filament diameter (0.338 nm/pixel). To maximize electron efficiency, we first restricted the effective imaging area of our Gatan K3 direct electron detector (5760 x 4092 pixels) to even dimensions in X and Y (4092 x 4092 pixels) (a step unnecessary for detectors with inherently square formats). Given the circular geometry of the electron beam and the square detector size, we optimized the beam diameter to match the camera dimensions at the chosen magnification (1.38 µm). This minimizes beam overlap and reduces unnecessary sample exposure without requiring modifications to standard Krios microscope settings, such as a square aperture. We selected tile overlaps of 10–15% (∼400-600 pixels) in both X and Y directions to ensure robust tile stitching. Using these parameters, a montage of a lamellipodium or cytoplasmic fragment would require dozens of overlapping tiles.

To achieve this, we implemented an acquisition strategy based on PACE-tomo (Eisenstein et al., 2023), which enables parallel tilt series acquisition using beam-image shift over user-defined polygonal regions in SerialEM (Mastronarde, 2005) (Figure 1A-1). Importantly, and particularly relevant for such large FOVs as acquired in this study, PACE-tomo incorporates a geometry-based sample model for improved tracking and autofocusing, and accounts for defocus variations with respect to the distance to the tilt axis. Using this acquisition strategy, we obtained multiple large-scale datasets, showing here two cells and one cytoplasmic fragment (Figure 1B-D, Table S2). These datasets contained 23 to 80 tiles, spanning areas from ∼37 to 127 µm², and were acquired within ∼78 minutes to ∼270 minutes, respectively.

Individual tomograms reconstructed from the acquired tilt series were of high-quality allowing to visualize individual actin filaments forming different architectures (Figure 1B, S2A-D). Within the tomograms we could also reliably identify additional molecular entities (Figure S2B) such as branch junctions, ribosomes, intermediate filaments, and clathrin-coated structures. Notably, even in areas of 4-fold beam overlap, where the cumulative electron dose reached ∼400 e⁻/Å², no observable beam damage could be observed.

### Large-scale optical flow-based 3D tile alignment and stitching

Established methods for creating a continuous, large field of view from overlapping tilt series stitch the individual 2D tiles at each tilt angle before reconstructing the combined 3D tomogram (Brown et al., 2026; Chua et al., 2024, 2026; Hylton et al., 2026; Peck et al., 2022; J. E. Yang et al., 2023, 2026). Although effective for small, regularly arranged montages, this approach becomes increasingly difficult for the large, irregular acquisitions shapes used here.

Hence, we developed an alternative strategy in which individual tomograms are first reconstructed and denoised independently and subsequently stitched in 3D to generate a single large montage volume. Conventional 3D stitching approaches (e.g., Blendmont within the IMOD package, BigStitcher (Hörl et al., 2019), or TeraStitcher (Bria & Iannello, 2012) primarily rely on rigid or locally constrained alignment based on cross-correlation and have limited capacity to accommodate non-linear deformations within the specimen, particularly when handling datasets comprising a large number of tiles. However, stitching of large volumetric datasets is a well-recognized challenge in connectomics (Tavakoli et al., 2025). Hence, we adopted an optical flow-based method called SOFIMA (Scalable Optical Flow-based Image Montaging and Alignment) (Januszewski et al., 2024), originally developed for stitching, alignment, and warping of large 3D connectomics microscopy datasets (see methods).

Despite the inherent challenges of our cryo-ET data - anisotropic resolution resulting from the missing wedge, low SNR due to dose limitations inherent to cryo-ET, and the dense and complex organization of actin networks - SOFIMA robustly stitched our datasets into single montage volumes (see Methods for optimized parameters) (Figure 1B-D). Specifically, we validated accurate tile overlap using manual and semi-automatic actin filament tracing approaches to confirm that individual filaments are seamlessly and faithfully connected across tile boundaries (Figure S3). In addition, visual inspection of the full montage volumes revealed long individual actin filaments reaching lengths of several micrometers and extending across multiple tiles (Figure S4A, B). We note that in few areas sample imperfections, e.g. due to ice contamination (Figure S5A, B) interfered with accurate stitching.

Together, these results demonstrate that our data acquisition and large-volume cryo-ET stitching strategy enables accurate reconstruction of large-scale actin networks up to 120 µm², effectively 2-3 times larger than published montage attempts, and supports reliable filament tracing across tile boundaries throughout the entire montage volume.

### F-actin parameters at the leading edge

We next set out to next use quantitative tools to determine the amount and spatial distribution of polymerized F-actin within the imaged leading-edge regions. For this we focused on the montages obtained from intact cells. Previous studies have attempted to estimate filamentous actin content at the cell leading edge using fluorescence microscopy, biochemical fractionation or semi-quantitative electron microscopy/tomography approaches (Aroush et al., 2017; G. Dimchev et al., 2021; Fäßler, Javoor, Datler, et al., 2023; Koestler et al., 2009; Mueller et al., 2017; Small et al., 1995; Svitkina et al., 1997). These studies provided important insights into actin dynamics and turnover but lacked the molecular resolution and spatial continuity required to directly quantify filament organization within intact three-dimensional actin networks. By contrast, actin filament vectorization across our large montage cryo-ET reconstructions now enables direct quantification of filamentous actin throughout entire lamellipodial volumes. The cumulative filament length within the analyzed datasets corresponded to 1.08 cm (Cell1, ∼81.6 µm^2^), 0.55 cm (Cell2 montage#1, ∼37.4 µm^2^), 0.87 cm (Cell2 montage#2, ∼51.6 µm^2^) and 0.93 cm (Cell2 montage#3, ∼55.4 µm^2^), reflecting total amounts of 3.98 million, 2.04 million, 3.23 million, and 3.44 million polymerized actin subunits, respectively (Table S3). To estimate the concentration of filamentous actin, we calculated the total number of actin monomers from the cumulative filament length assuming an axial rise of 2.7 nm per actin subunit and approximated the corresponding three-dimensional hull volume encompassing the actin network. Using this approach, we determined filamentous actin concentrations of ∼895–1036 µM (Table S3). These values are in good agreement with previous estimates obtained by quantitative fluorescence densitometry in zebrafish keratocyte fragments (1200±300 µM) (Aroush et al., 2017). However, the improved measurements of our data, due to its higher spatial and structural resolution, allows considerably greater confidence in the accuracy of data quantitation. Assuming there is approximately two-fold more actin in the soluble, diffusible pool compared to the polymerized F-actin network (Aroush et al., 2017), the total actin concentration can be estimated at up to 2-3 mM.

We next asked how actin filament lengths are distributed throughout the lamellipodial network. Early platinum replica electron microscopy and negative stain TEM studies suggested that lamellipodia contain filaments of heterogeneous and graded lengths, ranging from short, branched filaments near the leading edge to longer filaments extending deeper into the network (Small et al., 1995; Svitkina et al., 1997). In parallel, the nucleation-release model proposed that protrusive actin networks contain a large population of short, rapidly turning-over filaments generated by continuous nucleation, capping, and release from the leading edge (Mogilner & Edelstein-Keshet, 2002; Pollard & Borisy, 2003). Analyzing one montage (Figure 1B, area of 81.6 µm^2^ with a total filament count of 53,632) we find that lamellipodial actin filaments span a remarkably broad length distribution, ranging from 36 nm - corresponding to the shortest filament length detected under our vectorization parameters and equivalent to the crossover repeat distance or half-pitch length of an actin filament - up to ∼4.75 µm (Figure 2A, B). While the majority of filaments were shorter than 500 nm (∼91.6%), a sub-population of unexpectedly long filaments was also observed, with ∼2.2 % of all filaments exceeding 1 µm in length containing a fraction of ∼16.5% of the total F-actin (Figure 2B). Similar trends could be observed in the additionally analyzed montage datasets (Figure S6-8 A-B).

**Figure 2:**
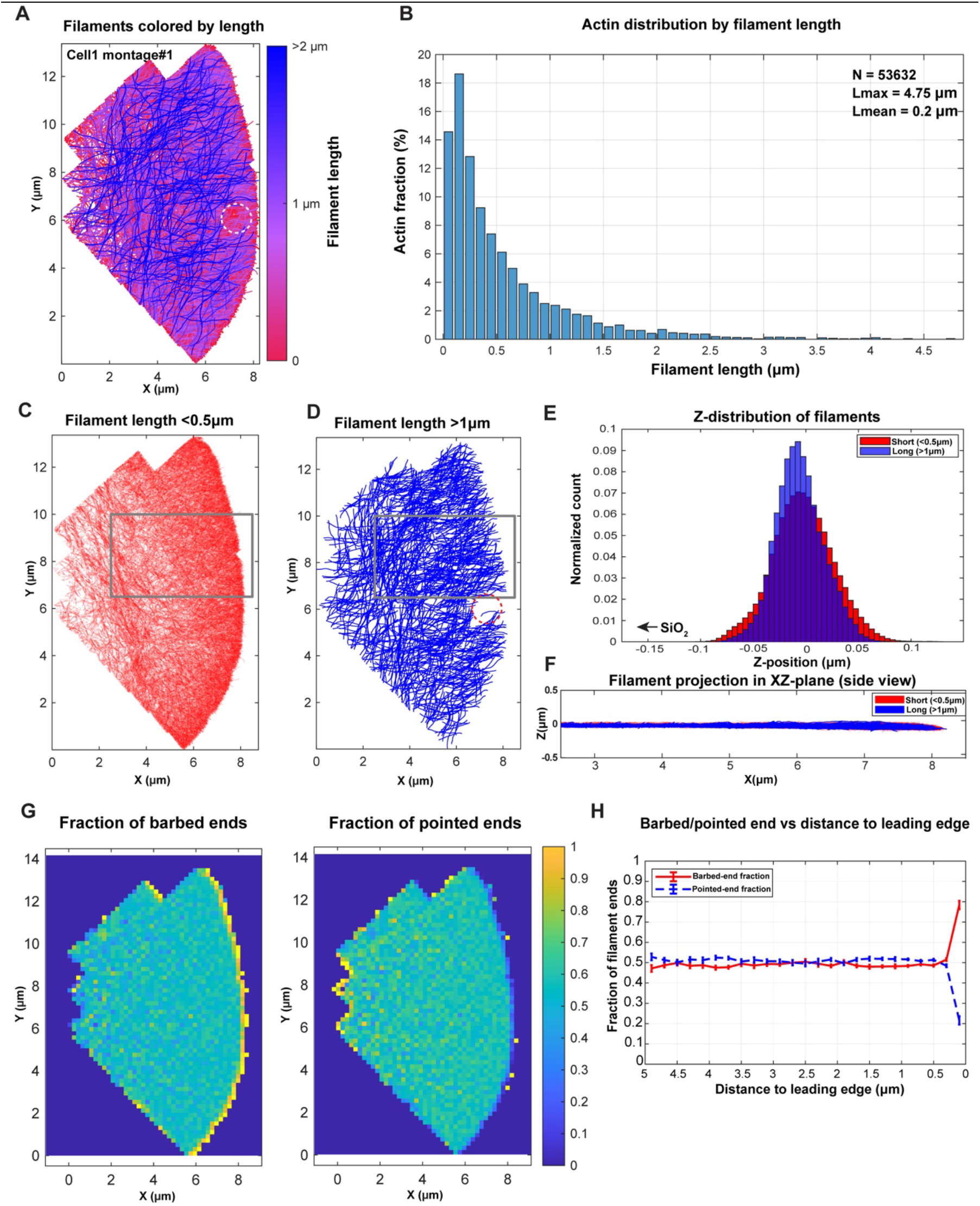
Quantitative analysis of actin filament network. **(A)** XY projection of all actin filaments traced in the cryo-CARE-denoised Cell1 montage #1, color-coded according to filament length. A region where filaments could not be traced properly due to ice contamination is marked by a dashed circle (also in D). **(B)** Actin distribution as a function of filament length, showing the fraction of total actin in each length bin. The total number of traced filaments (N), maximum filament length (Lmax), and mean filament length are indicated in the plot (Lmean). **(C, D)** Spatial distribution of filament-length subpopulations in the XY dimension. Short filaments, defined as filaments below 0.5 µm length, are shown in C, whereas long filaments, defined as filaments above 1 µm length, are shown in D. **(E)** Z-height position distribution of short filaments (<0.5 µm) and long filaments (>1 µm) (position of the SiO2 substrate is indicated for orientation). **(F)** XZ projection of short and long filaments from the boxed regions marked in C and D. **(G)** Spatial heat maps showing the local fraction of barbed ends and pointed ends across the montage. **(H)** Quantification of barbed-end and pointed-end fractions as a function of distance from the leading edge.

To further characterize the spatial organization of these filament populations, we mapped the localization of short (<500 nm) and long (>1 µm) filaments throughout the lamellipodial network. Short filaments were distributed across the entire protrusive network but displayed increased density near the leading edge. In contrast, long filaments were uniformly distributed throughout the network with filaments oriented parallel and perpendicular to the leading edge, suggesting long filaments might be involved in both force generation and in providing stability to the network, respectively (Figure 2C-D and S6-8 C-D).

We next investigated whether distinct actin filament populations segregate into separate layers within the lamellipodium, as proposed in a model invoking mechanically and dynamically distinct actin networks (Giannone et al., 2007). To address this, we extracted regions spanning from the leading-edge tip up to 4–5 µm into the cell interior and analyzed filament distributions along the z-axis. Cross-sectional analysis revealed that the actin network forms a continuous three-dimensional structure approximately ∼200 nm in thickness, in which short and long filament populations are extensively interwoven (Figure 2E, F). Notably, we did not observe evidence for two spatially segregated actin layers within the analyzed lamellipodial regions, in any of our analysed datasets (Figure S6-8 E-F).

Finally, the observation of a homogenous distribution of both long and short filaments throughout the entire depth of the network implies a balanced distribution of both barbed and pointed filament ends within the network. To address this, we plotted the density of barbed ends and pointed ends assuming that the filament ends more proximal to the membrane represent the barbed ends and the opposite the pointed ends. While the barbed end fraction is enriched in the first ∼300 nm from the leading edge as expected, we otherwise found a homogenous and equal distribution of ends up to 5 microns away from the tip of the lamellipodium into the rear of the network (Figure 2G-H, S6-8 G-H). Together, these analyses establish a quantitative framework linking nanoscale actin organization to mesoscale protrusion mechanics in migrating cells and reveal that long actin filaments extending several microns are a prominent and previously underappreciated component of lamellipodial architecture.

### Spatial heterogeneity of networks in the lamellipodium

Previous EM studies have linked lamellipodial actin architecture, including filament orientation and density, to protrusive activity and load adaptation (Koestler et al., 2008; Mueller et al., 2017). However, limited fields of view have made it difficult to follow network organization continuously from the leading edge to the rear of a lamellipodium or to capture long filaments in their entirety.

We therefore quantified filament orientation, density, and length across extended regions of motile keratocyte lamellipodia with locally convex, flat, and concave leading-edge geometries (Figure 3). Cell1 montage#1 displayed a broadly convex leading edge, consistent with a coherently advancing, protrusive lamellipodium. Within this apparently uniform protrusion, a dense population of predominantly short filaments roughly perpendicular to the leading edge progressively transitioned toward longer, more shallowly oriented filaments deeper in the network, becoming largely parallel to the leading edge toward the rear. The large field of view thus captures in 3D the continuous transition from a protrusive lamellipodial architecture toward a more lamella-like organization while preserving long filaments that would be truncated in smaller volumes (Figure 3A-C).

**Figure 3:**
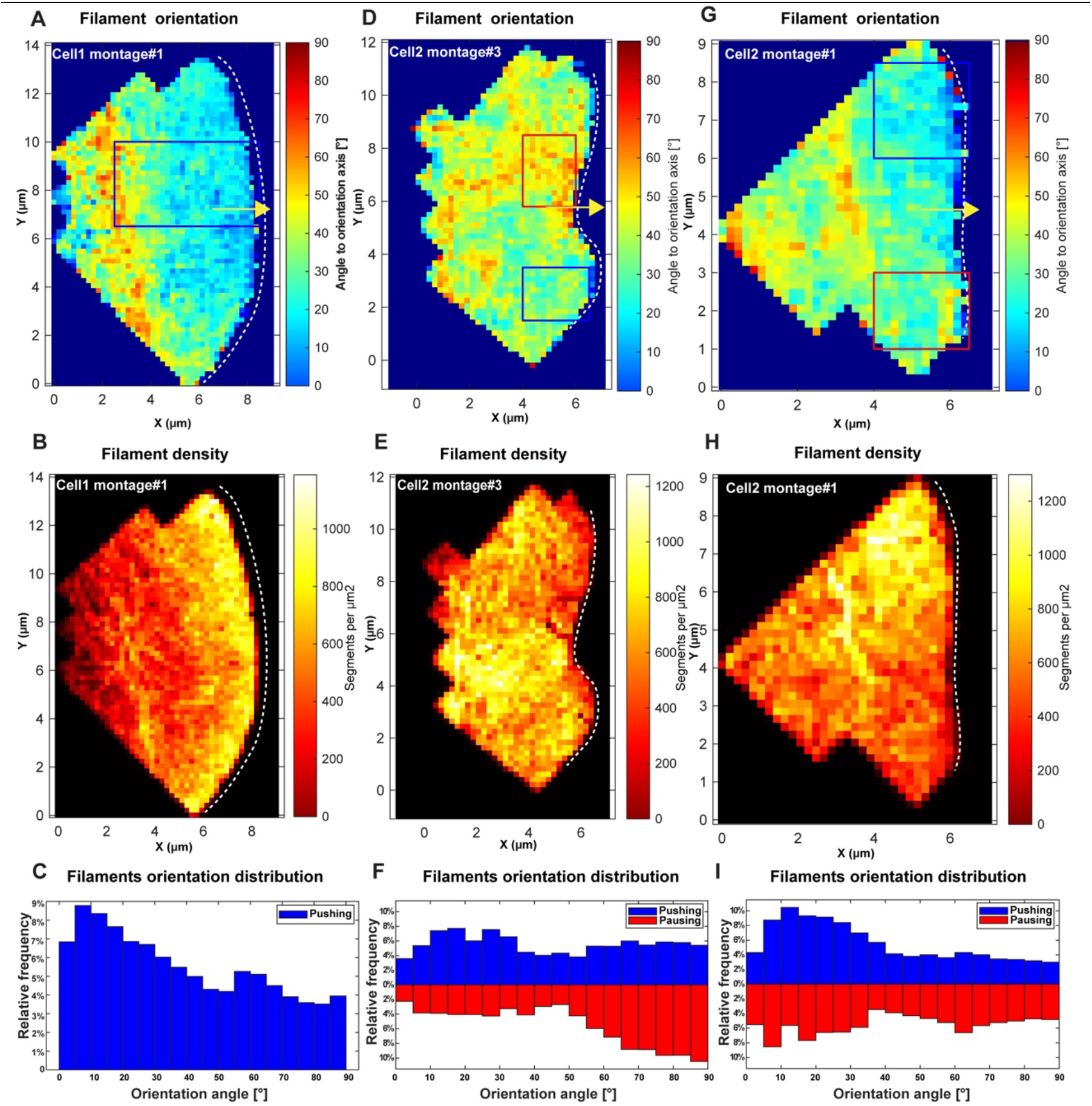
Local heterogeneity in actin-network topology. **(A, D, G)** Spatial maps of filament orientation calculated relative to the migration direction, indicated by the yellow arrow. The leading-edge shape is indicated by the white dashed lines. Low angles represent filaments oriented shallower to the migration direction and therefore more perpendicular to the leading edge, whereas high angles represent filaments oriented more transversely to the migration direction and therefore more parallel to the leading edge. Boxed regions indicate local regions used for the orientation-distribution analysis shown in C, F, and I. Blue boxes mark convex, protrusive-like regions, whereas red boxes mark flatter or concave regions with a more pausing-like morphology. Filament density and orientation were mapped in 2D using spatial binning. Density was calculated for each spatial bin (0.2 µm X 0.2 µm) as the number of filament segments divided by the bin area (segments per µm^2^)). Local orientation was calculated from the filament 3D end-to-end vectors within each bin, as the angle relative to the migration direction (yellow arrow). **(B, E, H)** Corresponding filament density maps for the same montage regions shown in A, D, G. **(C)** Filament-orientation distribution from the boxed region in A for Cell1 montage#1. **(F, I)** Filament-orientation distributions from the boxed protrusive and putatively pausing-like regions in D and G, respectively.

Cell2 montages #2 and #3 additionally revealed lateral variation along the same leading edge (Figure 3D-I). Regions with higher filament density and orientations biased toward the direction of protrusion alternated with regions containing a lower-density of longer filaments oriented more parallel to the leading edge, coinciding with local differences in membrane curvature. Thus, even within an apparently coherently advancing keratocyte lamellipodium, distinct local network architectures associated with different protrusive states can coexist along the leading edge.

Together, these analyses show how large-field-of-view cryo-ET captures both front-to-rear and lateral transitions in actin-network architecture, providing a continuous structural view of locally protrusive, pausing, and retracting states within a single migrating lamellipodium (Figure S9).

### Connectivity analysis of actin filaments within lamellipodial networks

Arp2/3-mediated nucleation at the plasma membrane tip of the leading edge is essential for the formation of lamellipodial actin networks (V. Dimchev et al., 2021; Fäßler, Javoor, Datler, et al., 2023; Suraneni et al., 2012; C. Wu et al., 2012).

To define the distribution of Arp2/3-mediated nucleation events at the level of single branch junctions, we used the molecular resolution of our montage cryo-ET data to identify positions of Arp2/3 branch junctions across the tomographic volume of montage#1 of cell2. For this we employed a refined template matching and classification approach (Figure S10, Methods), which allowed us to detect 6,954 candidate Arp2/3 branch junctions in the entire montage reconstruction of ∼37.4 µm^2^ (Figure 4A). To assess the reliability of branch detection, we extracted and manually insepected subtomograms centred on the identified branch junction positions. Approximately 92% of the detected candidates could be confirmed as bona fide branch junctions, corresponding to a false-positive rate of ∼8% (Figure S11A). This residual error is likely due to the filament crossings that mimic branch junction-like densities which are difficult to differentiate from true branch junctions within densely packed tomograms with anisotropic resolution and low signal to noise ratio (SNR). Conversely, template matching and stringent classification may fail to detect a fraction of true branch junctions, resulting in false negatives.

**Figure 4:**
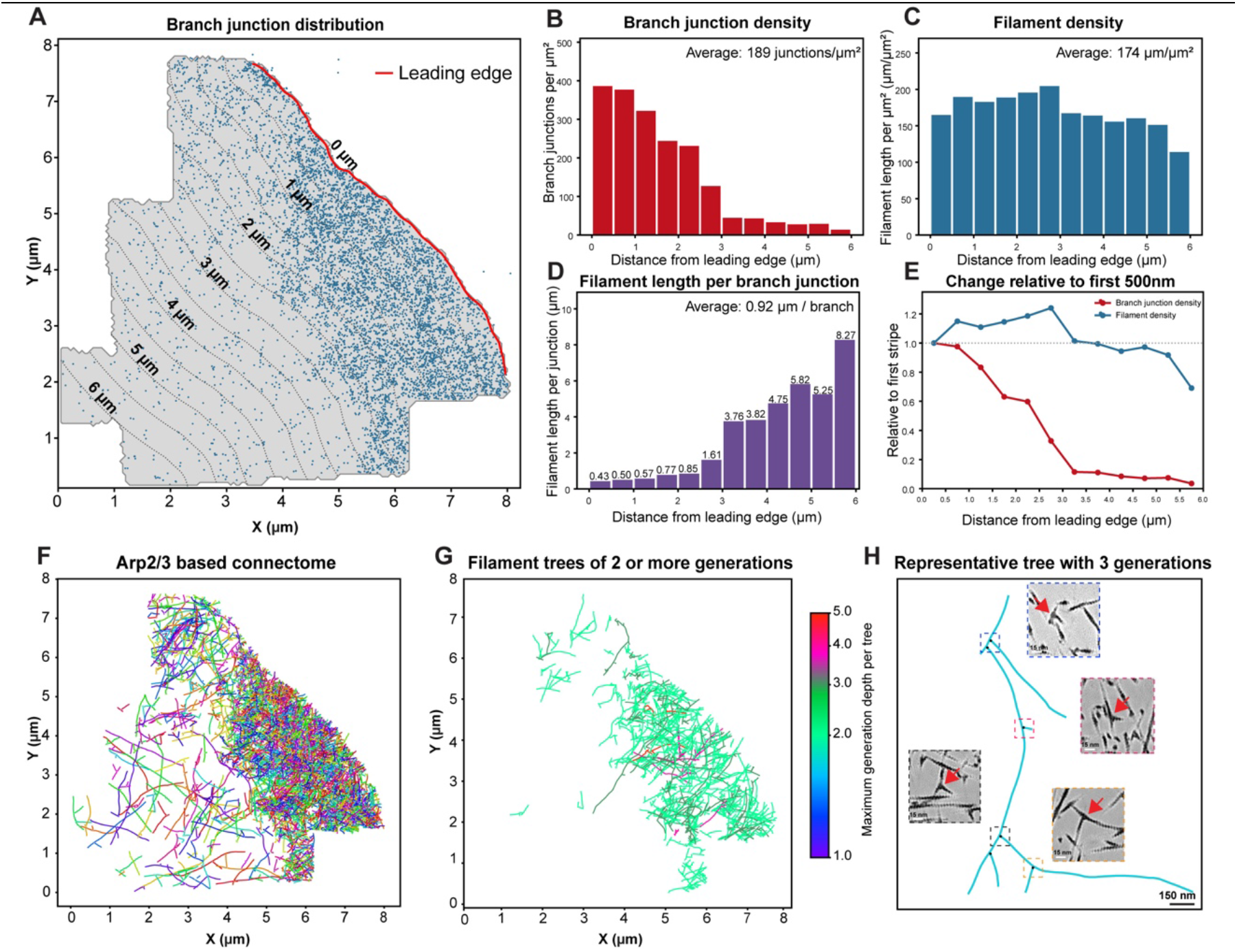
Distribution of Arp2/3 branch junctions and connectivity analysis. **(A)** Branch junction distribution in Cell2 montage#1. XY projection of all traced actin filaments in the IsoNet2 denoised montage was used to define the cell outline (grey). Template-matched branch junctions are shown as blue dots. The leading edge (red) was traced along the front of the filament outline, with the front direction taken as the side where the branch junction density ends abruptly. From the leading edge, the cell was divided into stripes of 500 nm width. Stripes up to 6 µm from the leading edge were used for the analysis in B-E. **(B)** Branch junction density, calculated as the number of branch junctions in a stripe divided by the area of the stripe. **(C)** Filament density, calculated as the total filament length in a stripe divided by the area of the stripe. **(D)** Filament length per branch junction, calculated as the total filament length in a stripe divided by the number of branch junctions in the same stripe (µm of filament per branch). **(E)** Change in branch junction density (red) and filament density (blue) relative to the first 500 nm stripe. **(F)** Actin filaments connected by Arp2/3 branch junctions were linked to reconstruct filament trees. The panel shows a 2D projection of differently colored Arp2/3-connected filament trees. **(G)** Arp2/3-branched filament trees with two or more generations of daughter filaments. Trees are color-coded by their maximum generation depth, as shown in the color bar. **(H)** Example filament tree with three generations of daughter filaments. Lines represent filaments and black dots mark the Arp2/3 branch junctions connecting them. Insets show the corresponding Arp2/3 branch junctions in the Isonet2 Cell2 montage#1 tomogram after the 3D tree model was overlaid onto it. Inset border colors match the boxed regions on the tree, and red arrows mark the junctions. Scale bars: 150 nm (main panel), 15 nm (insets).

Further inspection showed that many branches do not necessarily have long daughter filaments but rather resemble branch stumps, potentially the result of early termination of filament growth or daughter filament severing (Figure S11B). We found a subset of mother filaments with two branch junctions next to each other (double branches) (Figure S11C-D).

Thus, while the absolute number of detected branches may be affected by small errors in both directions, we expect the measured abundance to approximate the true branch-junction number, also based on earlier observations.

Our network contained ∼189 branch junctions per µm², corresponding to approximately one branch junction per 0.92 µm of filament contour length. This is remarkably close to earlier tomography estimates based on manual filament tracing and branch identification in mouse fibroblasts, which reported approximately one branch per ∼0.75–0.8 µm of filament length within a single tomogram of ∼ 1 µm^2^ area directly at the leading edge (Vinzenz et al., 2012). This agreement is notable given that our analysed montage reconstruction covers a substantially larger area and extends farther from the leading edge, where branch density is expected to decrease, as shown by previous light-microscopy and electron microscopy studies.

To confirm this, we calculated the density distribution of branches and actin filaments with respect to their distance to the leading edge. This showed that although there is a lateral variation in the branch density distribution (Figure 4 A), it does not seem to influence distance to leading edge distribution substantially and de-branching occurs faster than filament de-polymerization (Figure 4 A-C). Specifically, while within 1500nm from the leading-edge branch density remained rather steady (∼1 branch per 0.5 µm filament length), the branch density then quickly dropped to ∼1 branch every 4-8 µm filament length (Figure 4 D-E).

Together, these results demonstrate that template matching can be applied to montage tomograms to identify Arp2/3-dependent branch junctions with higher confidence than previous indirect measurement methods. While fluorescence studies reported parameters such as normalized Arp2/3 intensity (G. Dimchev et al., 2017; Fäßler, Javoor, Datler, et al., 2023; K. Lee et al., 2015), and small field-of-view electron tomography inferred Arp2/3/F-actin ratios (Vinzenz et al., 2012), our data now reports molecular-scale branch count over tens of square microns.

Importantly, modest systematic deviations in the absolute branch count are unlikely to substantially affect the relative spatial distribution of detected branch junctions, allowing us to robustly map local variations in branch density across the reconstructed lamellipodium.

Persistence of branch junctions away from the leading edge shows that not all the Arp2/3-mediated branches undergo debranching behind the membrane-proximal zone at the leading edge but remain embedded within the actin network as it matures and moves away from the leading edge due to retrograde flow. This observation raises a general question: do Arp2/3 branch junctions distant from the densely branched zone at the leading edge represent isolated branch remnants, or do they form connected branch architectures that have survived debranching and network turnover? To address this, we used the branch positions together with the traced filament coordinates to reconstruct 3D connectivity relationships between mother and daughter filaments. This allowed us to build branch-connected trees representing Arp2/3-mediated filament networks that remain physically connected within the tomographic volume (Figure 4 F, G, Figure S12A). To further validate actin network connectivity, we overlaid randomly selected reconstructed branch trees onto the raw tomographic volume. These overlays showed a clear Arp2/3 branch-junction density at each inferred mother-daughter connection, confirming that these connections reflect actual branch points in the network rather than tracing artifacts (Figure 4 H).

The resulting connectivity analysis further confirmed earlier data that the lamellipodial branch network cannot be considered as a single uniform branched entity (Vinzenz et al., 2012). Instead, multiple branch-connected trees are embedded within the dense actin meshwork (Figure 4 F, G). These trees varied substantially in size and topology, ranging from small local branch assemblies to larger connected structures generated by multiple successive rounds of branching yielding branch hierarchies with multiple generations of successive daughter-filaments (Figure S12 A, B).

With the large size of our montages we could follow long filaments to define how they are connected/embedded into the branched network. Interestingly, we identified numerous long filaments (even beyond 2 um length) throughout the lamellipodium without any detectable branches (Figure S13A). Specifically, the presence of these long unbranched filaments extending towards the very lamellipodium tip suggests that they may represent a distinct filament sub-population, potentially generated or maintained in a different way than the canonical branched network. These filaments may also be stabilized, allowing them to persist without rapid depolymerization. Interestingly, some of the long filaments (both that contained branches or not) displayed kinks or sharp bends, which have not been identified in previous EM studies (Figure S13A, B). Whether they represent sites of filament severing or the onset of depolymerization, debranching, or another not yet explained structural feature of actin networks remains to be elucidated.

## Discussion

In this work, we advance cryo-ET of cellular actin networks on two complementary fronts. Methodologically, we establish a large-scale montage cryo-ET workflow relying on optical flow-based 3D stitching, allowing extended field-of-view imaging with molecular-level structural information while maintaining compatibility with standard tomographic processing and downstream analysis strategies. Biologically, we provide, to our knowledge, the largest and most detailed three-dimensional visualization of the actin network in a protruding lamellipodium to date. This enables quantitative measurements of filament abundance, length distributions, spatial organization, branch-junction density, and network connectivity across lamellipodial length scales. Together, these data provide structural parameters that can inform the interpretation of previous functional and imaging studies, guide biophysical simulations, and generate new hypotheses about how lamellipodial actin networks are organized, remodeled, and mechanically integrated during cell migration. To facilitate further interrogation, we provide the tomograms and filament traces for download and through a public webKnossos-based web resource (see Data availability) that enables visualization of the filament network (Figure 5) (Boergens et al., 2017). This should support independent validation, reanalysis, and the development of computational models connecting molecular architecture to network mechanics.

**Figure 5:**
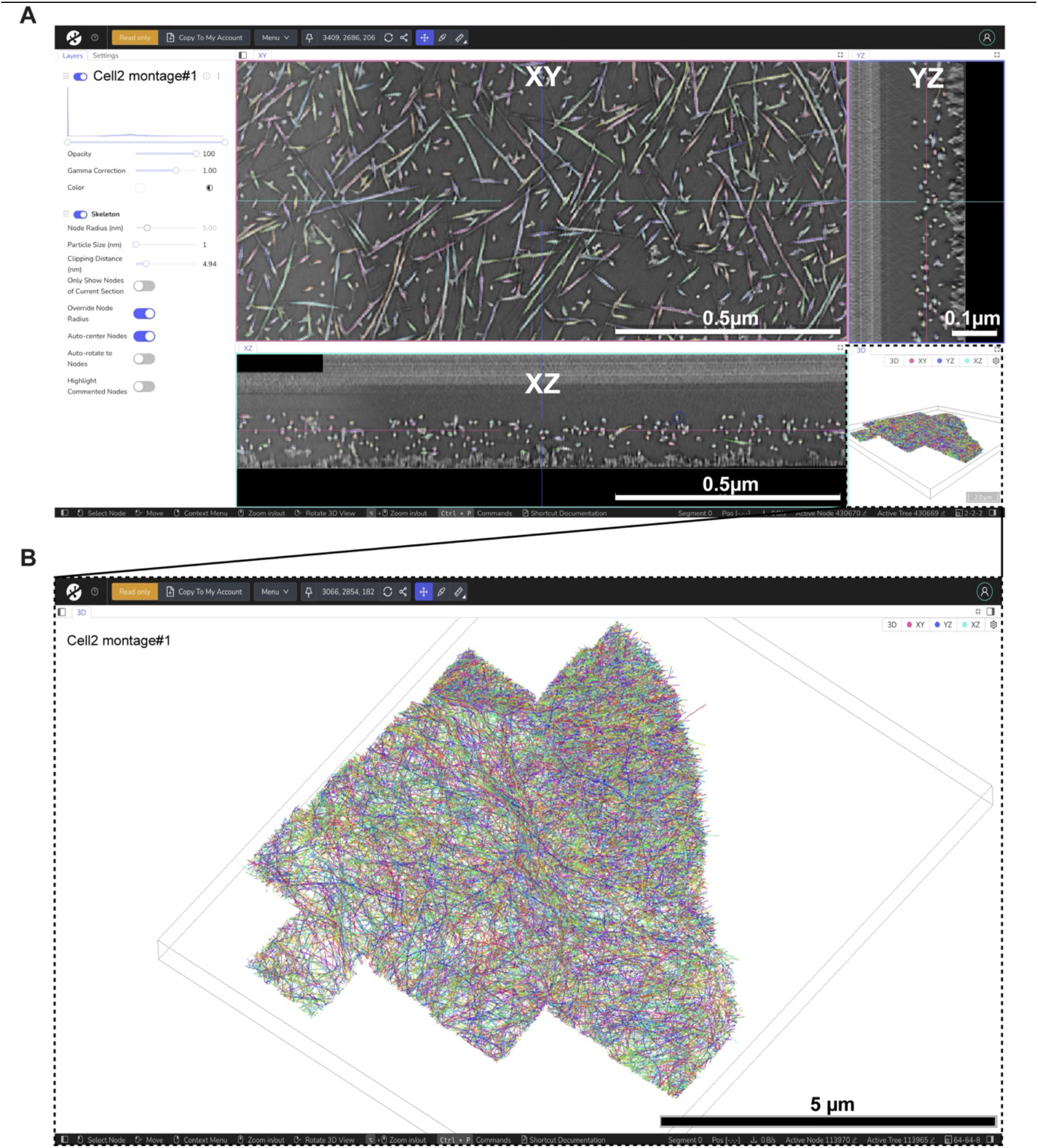
Public web-based access to stitched montage tomograms and actin filament traces. Stitched montage tomograms and their actin filaments traces are made available as a public, web-based resource through webKnossos (Boergens et al., 2017), allowing visualization and reanalysis. **(A)** Image of the webKnossos in-browser interface showing the IsoNet2 denoised Cell2 montage#1 overlaid with traced actin filaments. **(B)** Enlarged 3D view of the region boxed in A, showing all traced actin filaments of Cell2 montage#1, with each filament in a different color. Scale bars: 0.5 µm (XY and XZ views), 0.1 µm (YZ view), 5 µm (B).

### Implications for actin network understanding in lamellipodia

Our large-scale, molecular resolution data allow moving structural analysis of lamellipodia toward a connectomics-like framework, in which individual filaments and branch junctions are treated as elements of a spatially resolved network rather than as isolated features.

The measured F-actin concentration of approximately 0.8–1 mM, the total number of branches and their distribution, as well the distinction of filament populations based on connectivity provide important boundary conditions for existing models describing lamellipodia-mediated protrusion (Enculescu et al., 2010; Keren et al., 2008; J. Lee et al., 1993; Manhart & Schmeiser, 2017; Mogilner et al., 2020; Ponti et al., 2004, 2005; Watanabe & Mitchison, 2002; Weichsel et al., 2012). The agreement of our numerical observations with earlier fluorescence- and EM-based estimates for both actin concentration as well as overall branch junction density cross-validates the fidelity of our montage and the published indirect measurements (Aroush et al., 2017; Koestler et al., 2009), while our direct tracing now provides spatial and topological information unavailable from bulk or small field of view measurements.

The network is dominated numerically by short filaments but also contains long, frequently unbranched filaments extending over several micrometers. Rather than forming vertically segregated systems, these populations are interwoven into a composite network, refining earlier models proposing spatially and dynamically distinct lamellipodial and lamellar networks (Ponti et al., 2004, 2005; Vallotton & Small, 2009). One the one hand, short Arp2/3-dependent assemblies may support rapid, local force generation as well as maintaining the overall height of the lamellipodium. Here the double branches formed in close succession on the same mother filament (i.e. closer than the filament cross-over distance of 36 nm) could play a role, given the resulting variation of the angle of daughter filament growth with respect to the lamellipodium plane (Figure S11C-D) (Pollard & Borisy, 2003; Svitkina & Borisy, 1999). Long filaments could couple neighboring branch trees mechanically, sustain mechanical stress across the protrusion, preserve structural coherence during rapid turnover or eventually act as substrates for the long filaments embedded in the contractile network at the rear of the lamella. The origin of these long filaments remains unresolved: they could arise through formin-mediated nucleation/elongation - as suggested in studies that depleted FMNL in lamellipodia, Ena/VASP-dependent elongation, selective stabilization, or remodeling of initially branched filaments (Bear et al., 2002; Block et al., 2012; Kage et al., 2017; Rottner et al., 2017). Thus, their presence expands the canonical dendritic model by identifying an additional architectural component with potentially distinct dynamics and mechanics.

Mapping Arp2/3 junctions further shows that the lamellipodium is not a single continuous dendritic array but contains multiple branch-connected trees embedded within a dense filament meshwork, consistent with previous small field of view ultrastructural observations (Vinzenz et al., 2012). Regularly observed branch stumps may record rapid capping, or incomplete disassembly-processes implicated in lamellipodial network turnover (Blanchoin et al., 2014; Pollard & Borisy, 2003). Closely spaced double branch junctions may reflect spatial clustering of Arp2/3 activation rather than rapid sequential nucleation by a single WRC molecule. Because the catalytic reuse rate of individual WRC complexes has not been established, distinguishing repeated activity of one complex from activation by neighboring WRC molecules will require time-resolved single-molecule or reconstitution experiments, as well as spatially resolving the organization of the nucleation and elongation machinery on the membrane tip (Bieling & Rottner, 2023; Rottner et al., 2021; M. Wu et al., 2025).

The observed lateral and front-to-rear variations in filament density, length, orientation, and branch connectivity further shows that lamellipodial mechanical output is spatially patterned. Actin network force generation and load adaptation depend on filament density, length, orientation, and branching geometry (Bieling et al., 2016; Mueller et al., 2017; Schaus et al., 2007). In this regard it is interesting to visualize distinct architectures associated with different leading-edge geometries which coexist within one protrusion, indicating that apparently coherent migration can emerge from locally heterogeneous networks. Integrating these structural maps with correlative measurements of membrane dynamics and the localization of regulators such as WAVE, Arp2/3, formins, Ena/VASP, cofilin, and capping proteins should help establish how molecular regulation specifies local protrusive, paused, or retracting states. For this, combining the here shown montage cryo-ET approach with a method enabling temporal resolution, such as live-cell imaging would be particularly powerful (Auinger & Small, 2008; Inaba et al., 2024). Further work should also extend to other cell types displaying more diverse protrusive activities and filament architectures such as filopodia or microspikes that are absent in keratocytes.

Several limitations impact our interpretations. We study a system without membrane, not allowing to infer conclusions about membrane-filament interactions, such as would be needed to study actin filament nucleation and elongation at the very leading edge. Membrane removal might also lead to loss of short actin oligomers not strongly connected the network. Interruptions during tracing may underestimate filament length and connectivity, whereas some filaments or branch junctions may remain undetected in crowded regions or areas of lower SNR and stitching quality. Filaments leaving montage boundaries are necessarily truncated, and polarity inferred from membrane proximity is uncertain, particularly because pointed ends terminating at Arp2/3 junctions are not represented as free ends. Alternative approaches for polarity determination using structural features of individual filaments, combined with identified Arp2/3 complex branch junctions to define pointed ends could be implemented (Burbaum et al., 2021; Martins et al., 2021). Nevertheless, the recovery of continuous micron-scale filaments and reproducible network-scale trends indicates that these limitations do not impact the principal conclusions. Together, the data support a view of the lamellipodium as a crowded, heterogeneous composite of transient branch-connected modules and persistent long filaments whose integration generates protrusion at the cellular scale.

### Optical flow-based 3D stitching for large field-of-view cryo-ET

A central feature of our workflow is that individual tomographic tiles are reconstructed independently and subsequently stitched in three dimensions. This differs from previous montage strategies that align two-dimensional projections before reconstruction (Brown et al., 2026; Chua et al., 2024, 2026; Hylton et al., 2026; Peck et al., 2022; J. E. Yang et al., 2023, 2026). While projection-based stitching can be effective for smaller or regular montages, errors from local deformation, imperfect geometry, or alignment inaccuracies may propagate into the final volume. By stitching reconstructed tomograms in 3D, our approach preserves local reconstruction quality and enables non-rigid correction of tile-to- tile distortions directly in volume space.

The dense lamellipodial actin network provides a stringent test case for this strategy. Actin filaments are only a few nanometers wide, densely packed, and embedded in low-contrast cryo-ET data with anisotropic resolution. At the same time, filament continuity across tile boundaries offers a direct biological metric for stitching accuracy, something that has not yet been done for other published montage workflows. The seamless tracing of filaments over tile boundaries therefore validates the alignment not only geometrically, but also at the level of the molecular structures relevant for downstream analysis.

An important advantage of reconstructing tiles independently is modularity. Individual tomograms remain available for structural analysis on raw data, as well as denoising of the individual tiles, a problem recently recognized for large montages reconstructed from stitched 2D-tiles.

The stitched, denoised montage volume provides the spatial context required for network-scale interpretation. Thus, the montage does not strictly need to enable high-resolution analysis. It rather links molecular information to cellular-scale organization. This is particularly relevant for visual proteomics, where molecular abundance, localization, and spatial relationships must be interpreted across extended cellular regions.

A practical requirement of montage cryo-ET is overlap between neighboring tiles, necessary for robust registration. We found 10-15% overlap to work well, similar to what has been recently shown as also being required in other montage acquisition schemes (Brown et al., 2026; Chua et al., 2026; Hylton et al., 2026; J. E. Yang et al., 2023, 2026). Consequently, local regions can receive increased cumulative dose. In our samples, even areas with the highest overlap did not show obvious beam-induced damage. However, dose tolerance is sample-dependent and should be optimized for each application. Still, other recent montage applications used high-doses on a variety of samples, showing that this is not necessarily an issue. Future improvements in beam shaping, detector performance, phase-plate technology (Petrov et al., 2026), and denoising are likely to further increase the efficiency and applicability of large-scale montage cryo-ET.

Hence, although our workflow developed here is applied to study lamellipodial actin networks, we expect it to be broadly applicable to biological systems whose relevant architecture extends beyond the field of view of conventional cryo-ET. Samples containing continuous, high-contrast features spanning tile boundaries, such as cytoskeletal filaments, membranes, cell–cell interfaces, bacterial envelopes, or organelle networks, are expected to be particularly well suited. More homogeneous or weakly contrasted samples may be more challenging but could benefit from additional fiducial markers to support registration.

## Data availability

The fish keratocyte F-actin structure and model are deposited under accession codes EMD-100760 and PDB 35PI, respectively. One montage tomogram and its corresponding filament vectorizations are available as webKnossos-based web links:

Cell2 montage#1: https://webknossos.org/links/p6_q1V2brvf8vN55

## Code availability

The custom scripts used for data processing and analysis will be made publicly available after the final acceptance of the manuscript.

## Material availability

This study did not generate new unique reagents.

## Lead contact

Requests for further information and resources should be directed to and will be fulfilled by the lead contact, Florian Schur

## Acknowledgements

We would like to thank the group of Johann Danzl at ISTA for their initial input on using SOFIMA for stitching 3D volumes. We also thank the C.P. Heisenberg group and Verena Mayer from the fish facility at ISTA for providing zebrafish for our experiments. This research was supported by the Scientific Service Units (SSUs) of ISTA, including resources provided by Scientific Computing (SciComp), the Life Science Facility (LSF), the Imaging and Optics Facility (IOF), and the Electron Microscopy Facility (EMF). We acknowledge funding from ISTA, the European Research Council (ERC) Starting Grant (European Research Council (*ERC*) of the European Union grant ActinID, No. 101076260) awarded to F.K.M.S., and the ERC Synergy grant (PushingCell, No. 101071793) awarded to M.S. Views and opinions expressed are however those of the author(s) only and do not necessarily reflect those of the European Union or the European Research Council Executive Agency. Neither the European Union nor the granting authority can be held responsible for them.

## Author contributions

Conceptualization: MGJ, FKMS; Methodology: MGJ, VVH, CS, RH; Investigation and formal analysis: MGJ, K.v.d.E, CS, RH, VVH; Data curation: MGJ, FKMS; Visualization: MGJ, RH, K.v.d.E, CS, FKMS; Software: MGJ, K.v.d.E, CS, RH, VVH, BS; Validation: MGJ, K.v.d.E, NW, FKMS; Writing-original draft: MGJ, FKMS; Writing-review & editing: MGJ, K.v.d.E, CS, RH, VVH, MS, FKMS; Project administration: MGJ, FKMS; Resources: FKMS, MS; Supervision: FKMS, MS; Funding acquisition: FKMS, MS.

## Declaration of generative AI and AI-assisted technologies in the writing process

During the preparation of this work, the authors used ChatGPT (OpenAI), Claude (Anthropic), and Gemini (Google) in order to assist with script writing and for consolidating own scripts used for data analysis and for literature search and improving the clarity of the text. After using these tools, the authors reviewed and edited the content as needed and take full responsibility for the content of the published article.

## Materials and Methods

### Preparation of keratocytes and keratocyte fragments

Wild-type zebrafish (approximately 2 years old) were maintained at 28 °C in the zebrafish facility at ISTA. Keratocytes were prepared from scale explants using a previously published protocol with minor modifications (Mueller et al., 2017; Small et al., 1995; Sun et al., 2016). Fish were euthanized according to institutional guidelines, and approximately 30-40 scales were removed from each fish irrespective of sex. Scales were washed three times in DMEM (Gibco) and transferred into 6-well plate containing 1 mL START medium per well, with 15-20 scales per well. START medium comprised 70% fish Ringer, 20% DMEM, and 10% Steinberg medium supplemented with 10 mM PIPES (pH 7.5) and 2% chicken serum. Fish Ringer consisted of 112 mM NaCl, 2 mM KCl, 2.4 mM NaHCO_3_, 1 mM CaCl_2_, and 1 mM Tris (pH 7.3), whereas Steinberg medium consisted of 52 mM NaCl, 0.3 mM Ca (NO_3_) ·4H_2_O, 0.6 mM KCl, and 0.8 mM MgSO_4_·7H_2_O. A sterile coverslip was placed over the scales, and a small weight (nut) was used to keep them gently flattened beneath the coverslip. Plates were incubated overnight at 28 °C to allow keratocytes to migrate out from the scales onto the surface of the dish. The following day, the scales were gently removed with forceps and migrated epithelial cells were washed three times with PBS and incubated for 40 min in RUNNING medium supplemented with 1 mM EGTA. RUNNING medium comprised 90% fish Ringer and 10% Steinberg medium supplemented with 10 mM PIPES (pH 7.0). Cells were then washed twice with PBS and treated with 0.25% trypsin-EDTA. After 1 minute, trypsin inhibitor was added at an equal volume to terminate the reaction. The resulting single-cell suspension of keratocytes was transferred onto glass coverslips or cryo-EM grids for further processing.

### Immunofluorescence microscopy of keratocyte cells and fragments

Coverslips were coated with fibronectin (25 µg/mL) for 1 hour at room temperature. After coating, the coverslips were washed three times with PBS and transferred into a 12-well plate. Approximately 300-400 freshly trypsinized keratocytes were seeded into each well and incubated in START medium at 28 °C for 1 hour to allow the cells to adhere and spread on the fibronectin-coated coverslips.

After 1 hour, for generation of cytoplasmic fragments the cells were treated with START medium containing 200nM Staurosporine for 30 min. Following this, the Staurosporine containing medium was replaced with START medium without Staurosporine and incubated for 1 to 2 hours at 28 °C to allow cells and fragments to adhere and spread.

After 1 hour of incubation at 28 °C, the medium was aspirated, and the cells and fragments were fixed with 4% paraformaldehyde (Merck, P6148) in PBS, pre-warmed to 28 °C, for 20 minutes. Following fixation, the coverslips were washed twice with PBS and permeabilized with 0.1% Triton X-100 in PBS for 60 seconds. After permeabilization, they were washed twice again with PBS.

The coverslips were then incubated for 60 minutes on 50 µL droplets containing 1 µg/mL phalloidin-488 at room temperature. After incubation, excess phalloidin was removed by washing twice with PBS. The coverslips were then mounted onto glass slides using mounting medium containing DAPI (Fluoromont-G). Slides were stored overnight at room temperature, then transferred to 4 °C for longer-term storage until imaging. The slides were imaged on a Zeiss Axio Imager (Zeiss) equipped with a CoolLED p300 SB light source (CoolLED) and an LCI Plan-Neofluar 63x/1.3 Imm Corr DIC M27 objective (Zeiss). Images were processed in Fiji for generating merges/overlays.

### Cryo-EM sample preparation of keratocyte cells and fragments

Trypsinized keratocytes were seeded onto Quantifoil SiO₂ R2/2 200 mesh or SiO₂ R1.2/20 150 mesh grids (Quantifoil Micro Tools). Prior to cell seeding, the grids were glow-discharged in a Quorum GloQube (25 mA for 90 seconds). The grids were then placed onto 3D-printed grid holders (Fäßler, Zens, et al., 2020) and coated with fibronectin (Sigma RH, 25 µg/mL) for one hour at room temperature. After coating, the grids were washed twice with PBS and transferred into a 96-well plate. Between 100 and 200 trypsinized keratocytes in START medium were seeded onto each fibronectin-coated grid. The cells were allowed to adhere and spread on the grids for one hour at 28 °C. For generation of cytoplasmic fragments, the START medium was replaced with START medium containing 200 nM Staurosporine and incubated for 30 minutes at 28 °C. Following this, the Staurosporine containing medium was removed, and the grids were rinsed with 100 µL of START medium (without Staurosporine). The grids were then incubated for an additional 1 to 2 hours at 28 °C to allow fragments to adhere and spread.

Prior to plunge freezing, the grids were subsequently removed from the grid holders and placed into 50 µL drops of cytoskeleton buffer (10 mM MES, 150 mM NaCl, 5 mM EGTA, 5 mM glucose, and 5 mM MgCl₂, pH 6.2) supplemented with 0.75% Triton X-100, 0.25% glutaraldehyde (Electron Microscopy Sciences, E16220), and 0.1 µg/mL phalloidin (Merck, P2141) for 50-60s at room temperature. The grids were then transferred to 50 µL drops of cytoskeleton buffer containing 2% (w/v) glutaraldehyde and 1 µg/mL phalloidin for 15 minutes at room temperature. The grids were vitrified using a Leica GP2 plunger (Leica Microsystems) under blotting chamber conditions of 90% humidity and 4 °C. Prior to vitrification, 3uL of PBS was applied to the grids and the grids were backside blotted for 3-4 seconds (SiO₂ R2/2 200 mesh) or 6-8 seconds (SiO₂ R1.2/20 150 mesh) using the Leica GP2’s internal blotting sensor, then plunge frozen in liquid ethane at -185 °C. The vitrified grids were stored in liquid nitrogen until imaging.

### Cryo-EM data collection

A single-particle cryo-EM dataset was acquired on keratocytes and fragments migrating on Quantifoil SiO₂ R2/2 200 mesh grids (see above for sample preparation) using a Glacios TEM microscope operating at 200 kV (Thermo Fisher Scientific) equipped with Falcon4i detector. EPU version 3.7.0 (Thermo Fisher Scientific) was used for automated hole selection and data collection. The raw 4096 x 4096-pixel movies were recorded at a nominal magnification of 150,000x resulting in a pixel size of 0.9551 Å. The Falcon4i direct electron detector was used in electron-counting mode, and the movies were saved in EER (Electron Event Representation) format. The defocus range was set to -0.5 to -3.0 µm and the cumulative dose was set to 80 e⁻/Å².

### Single-particle image processing

Single-particle image processing was performed in cryoSPARC v 4.4 (Punjani et al., 2017) (Figure S1D-F). The raw EER movies of 4096 x 4096 pixels were imported into cryoSPARC. Patch based motion correction and CTF estimation were performed using the default patch size in cryoSPARC. Micrographs were manually curated to remove empty micrographs or micrographs containing thick ice, resulting in a final set of 994 micrographs from 2424 originally acquired micrographs. Initial particle picking was performed using filament tracer with minimum diameter and maximum diameter set to 3 and 13 nm respectively. One round of 2D classification was performed on particles extracted with a box size of 360 pixels. Selected 2D classes of actin filaments were used as a template for subsequent template picking. After template picking, 386,378 particles were extracted with a box size of 360 pixels and subject to two rounds of 2D classification resulting in 17 good classes with 204,916 particles. Using these particles, ab-initio reconstruction and homogeneous refinement were performed with C1 symmetry. Further, helical refinement was performed using a twist and rise of -167 ° and 28 Å, respectively. After one round of 3D classification, based on map quality, one class containing 129,656 particles was subjected to helical refinement, local refinement and local CTF estimation. A final round of helical refinement on 129,656 particles resulted in 3.61 Å resolution map measured at the 0.143 Fourier shell correlation (FSC) criterion. The helical parameters of the final map displayed a helical rise of 28.33 Å and twist of -167.09 °.

### Model-building and real space refinement

For visualization of all EM-density maps and models, UCSF ChimeraX version 1.10.1 was used (Meng et al., 2023; Pettersen et al., 2021). Model building and refinement were performed in ISOLDE plugin version 1.10.1 (Croll, 2018) in ChimeraX.

To build a model of the zebrafish actin filament, we used an AlphaFold2-predicted model of zebrafish actin (Uniprot: A8WG05) from the Google DeepMind dataset (AF-A8WG05-F1-v6) (Jumper et al., 2021). This monomer model was rigid body-fitted into the EM density of the actin filament structure obtained from SPA. We then manually fitted ADP into visible densities based on an existing model of actin. As we only observed electron microscopy density for residues 6 to 372, we removed the remaining residues (1-5, 373-375).

To generate a filament model, we then rigid body-fitted 4 neighboring copies of the Actin-ADP monomer into the EM-density of the filament resulting in a total of 5 monomers occupying all the contact sites of the central monomer. Four noncrystallographic symmetry (NCS) copies of the monomers were restrained to the centrally positioned monomer that was iteratively refined locally in ISOLDE. Finally, the NCS copies were replaced with the refined, centrally positioned monomer and phalloidin molecules were rigid-body fitted in ChimeraX. One round of real-space refinement of the F-actin model was then performed in Phenix version 2.1-6048 (Afonine et al., 2018) using NCS constraints to ensure that all subunits and inter-subunits are identical resulting in the final model of F-actin.

### Comparison of the F-actin models

To compare the zebrafish F-actin model with a previously deposited actin filament structure, one monomer from the final zebrafish model was aligned with one corresponding monomer from phalloidin-bound chicken skeletal F-actin (PDB 7BTI) in ChimeraX. For this comparison, both monomers were restricted to the common fitted residue range (6– 372) and the C-alpha RMSD value was calculated from the aligned residues.

### Large-Scale cryo-ET montage data acquisition, processing and analysis

A graphical summary of the experimental steps described below is shown in Figure S14.

### Montage Cryo-Electron Tomography (Cryo-ET) data collection

Tilt series were acquired under cryogenic conditions on a Thermo Fisher Scientific Krios G3i transmission electron microscope (300 kV) equipped with a BioQuantum post-column energy filter and a K3 camera (Gatan), with fringe-free illumination. Gain reference images were collected prior to data acquisition. DigitalMicrograph 3.52.392.0 (Gatan) was used for Zero-Loss Peak (ZLP) tuning and with an energy filter of 20-eV slit width. Microscope alignment, astigmatism corrections, and data acquisition were performed using SerialEM (v. 4.1.10)(Mastronarde, 2005).

Low- and medium-magnification maps of cells and fragments were used to define areas of interest for subsequent high-resolution montage tomography acquisition. The electron beam diameter was reduced to fit the short axis of the K3 camera field of view, enabling the collection of square images (4092 × 4092 pixels) in x and y dimensions. This reduction is essential to avoid beam damage in overlapping exposure regions during montage acquisition, which would be increased due to the otherwise rectangular size of the K3 chip. A custom script in SerialEM was used to distribute a square grid of acquisition points over the selected area (SA map) of cells and fragments, maintaining a 10%-15% overlap between adjacent tiles (in both X and Y). Tilt series were acquired using the PACE-tomo (v. 1.4.4) (Eisenstein et al., 2023) workflow with a dose-symmetric tilt scheme ranging from -50° to +50° in 2° increments. The nominal defocus was set to -6 µm, and the nominal magnification was 26,000×, corresponding to a pixel size of 3.371 Å. Each tilt image was recorded as a movie of eight frames at 4092 × 4092 pixels. The dose per tilt was set to 1.76 and 1.96 e⁻/Å², resulting in a cumulative dose of approximately 90 and 100 e⁻/Å² per montage tilt series respectively (see table S2 for details). At the corner overlap regions, the highest tolerable dose (without inducing visible) sample damage was 400 e⁻/Å² using SiO_2_ support film grids which is higher than the carbon support film grids.

### Tomogram reconstruction and denoising

Tomograms were differently denoised depending on downstream processing and analysis. On the one hand, individual tilt series were motion corrected (5×5 patch size), followed by tilt-series alignment and tomogram reconstruction using the weighted-back projection (WBP) method as implemented in AreTomo3 (v. 1.3.4) (Peck et al., 2026; Zheng et al., 2022), resulting in 4x binned (pixel size = 13.50 Å) tomograms. To enhance contrast tomograms were first cropped in Z to remove empty slices (Z from 250 to 220) and denoised using cryo-CARE (v. 0.3.0) (Buchholz et al., 2018). Default parameters were used on even and odd pair raw tomograms (generated from even and odd frames) to obtain denoised full tomograms.

Alternatively, individual tilt series were pre-processed in WarpTools v2.0.0 (Tegunov & Cramer, 2019), followed by initial tilt series alignment in AreTomo2 v1.1.3 (Zheng et al., 2022) using patch tracking. These tilt series alignments were refined using Miss-alignment v0.2.0 (Chaillet et al., 2026) and refined alignments were then used to reconstruct 4x binned (pixel size = 13.50 Å) tomograms within WarpTools. Even/odd pair tomograms reconstructed from even/odd frames were subjected to missing wedge correction and denoising using IsoNet2 (Liu et al., 2025) to obtain denoised full tomograms.

Differently denoised tomograms were used as indicated in the methods sections and the figure legends showing the respective data.

### Pre-processing of individual tomograms for stitching

The pre-processing of cryo-CARE or IsoNet2 denoised tomograms for stitching into montage reconstruction involves several steps. Initially, each stack is rotated 90° counterclockwise and inverted using an ImageJ (Schindelin et al., 2012) script (required for downstream 3D stitching). The histogram of each of the volumes is then adjusted and normalized to standardize intensity values (0.2%, 99.8%) in MATLAB, and subsequently converted into a 16-bit data format. Location information for the individual tomograms (stage coordinates from PACE-tomo acquisition) and numbering are formatted for compatibility with BigStitcher (Hörl et al., 2019). In BigStitcher, stage coordinates are assigned to each tile for further reconstruction of the dataset. Finally, the entire dataset is exported in N5 format.

### Montage tomogram generation using SOFIMA

For stitching the individual tomograms into a single large-volume reconstruction, we used SOFIMA (Januszewski et al., 2024) (https://github.com/google-research/sofima), following and adapting the procedure previously described for stitching expansion microscopy datasets (Tavakoli et al., 2025).

Each 3D tomogram (tile) was preprocessed using contrast-limited adaptive histogram equalization (CLAHE) (Zuiderveld, 1994), applied independently to every xy-plane of the tile volume. Unlike the normalization step performed prior to BigStitcher, CLAHE adjusts local contrast non-linearly to enhance local structural feature visibility rather than global scaling of voxel intensities. The procedure was implemented with the *equalize_adapthist* function from the *skimage.exposure* module (scikit-image, version 0.25.2) using a clip limit of 0.02. The resulting intensity values (ranging from 0 to 1) were then scaled by a factor of 255 and converted to the 8-bit pixel type. Throughout the workflow I/O operations were handled using the *tensorstore* (version 0.1.76).

The tiles had a lateral plane size of 1022×1022 voxels and axial extent varying depending on experiment with expected lateral overlap of 10% or 15%. We then used a coarse positioning step to find their initial location in the coordinate system of the output volume, followed by fine alignment to elastically deform them to obtain seamless matches between adjacent tiles. The tiles were then rendered with linear distance-weighted blending of overlapping image content.

In the initial coarse positioning step, we isolated a cubic segment measuring 50 voxels from the ‘preceding’ tile within each adjacent tile pair. Utilizing cross-correlation techniques, we located the analogous image content within the ‘following’ tile, thereby establishing an offset vector representing the spatial displacement between the tiles. Subsequently, the tiles were conceptualized as unit masses linked by springs, with the springs’ resting lengths equivalent to the computed offsets. This configuration was allowed to reach a state of equilibrium, with the equilibrium state serving to establish the initial tile positions for the subsequent fine alignment phase.

In the fine alignment stage, each tile was represented as an elastic mesh composed of unit masses organized on a regular grid with a three-dimensional stride of 20 voxels. The nearest neighbor masses were interconnected by Hookean springs. Following coarse positioning, adjacent tiles were positioned to partially overlap. For each mass situated in these overlapping regions, cross-correlation techniques were employed to determine the location within the neighboring tile for a 50-voxel image patch centered on the mass. These procedures established point correspondences between the tiles, which were subsequently modeled as zero-length Hookean springs. The system was allowed to reach equilibrium, and the relaxed configuration of the tile meshes was saved and utilized to render the stitched volume.

#### Detailed SOFIMA parameters

During the rigid, coarse alignment phase, we established the relative configuration of tile pairs by utilizing the predefined tile layout corresponding to the image acquisition process. The parameters QUERY_R_ORTHO and QUERY_R_OVERLAP was set to a value of 40 voxel. The minimum overlap for tiles (QUERY_OVERLAP_OFFSET) was specified as 0, while the dimensions of the search area within tiles (SEARCH_OVERLAP and SEARCH_R_ORTHO) were defined as 350 and100 voxels respectively. The subsequent fine stitching step involved warping the tiles to facilitate a smooth transition at the borders, employing a voxel stride of (20, 20, 20) in the x, y, and z directions. This step produced a mesh that ensured globally consistent warping and maintained the relative arrangement of the individual tiles.

#### Rendering output

In the last step, we applied the obtained mesh to the cryo-CARE or IsoNet2 denoised tomograms for downstream actin filament tracing and raw tomograms for template matching, resulting in the respective stitched volumes rendered into the 8-bit MRC file format.

#### Computing requirements

We performed these processing steps on ISTA’s high performance computing cluster. Typically, for CLAHE, we requested 32 CPUs (Intel Xeon CPU E5–2680 v4 @ 2.40-GHz or similar) and 128 GB of RAM. For coarse and fine stitching, we typically requested 1 GPU (GeForce L40 with 48 GB of VRAM), 32 CPUs and 128 GB of RAM. Rendering was typically done requesting 32 CPUs and up to 256 GB of RAM.

### Validation of tomogram stitching and filament tracing

First, we randomly selected sub-volumes located at corner overlap regions from three stitched cryo-CARE montage tomograms, where four tomograms meet. These regions are particularly stringent tests because successful tracing must extend beyond the nominal 10-15% tile overlap. From 3 montage tomograms (cell2 montage#1-#3) acquired with 10% overlap (∼102 pixels (∼138 nm)), we cropped six sub-volumes of size 400×400x512 voxels (∼540nm x 540nm x 692nm) in IMOD (Kremer et al., 1996). Three independent expert annotators manually traced actin filaments in IMOD for all sub-volumes. In each case, experts were able to trace filaments continuously from one tile into the adjacent tile(s) running beyond the overlap regions, indicating that the stitched volumes preserved filament continuity across boundaries. In parallel, we used the Amira 2020.2 filament-tracing module to generate automated filament vectorizations for the same sub-volumes (see section “Vectorization of actin filament network”). We then compared manual and automated filament traces (Figure S3). A direct quantitative similarity metric (e.g., cross-correlation or difference maps) is not straightforward for filament traces, because point placement, particularly along the Z axis varies between annotators in densely packed regions. Nevertheless, visual comparison showed strong agreement between manual and automated filament traces, and long filaments spanning tile boundaries could be confidently identified by both methods. Together, these results indicate that the 3D stitching quality allows for faithful filament tracing across tiles.

Second, we leveraged the presence of long filaments that traverse multiple tiles in the complete montage tomograms to validate accurate tile stitching. One trained expert manually traced 10-12 long (up to 4.5 microns) filaments per montage running across multiple tiles, further supporting that SOFIMA stitching preserves filament network continuity (Figure S4). While we can trace long filaments, our montage volumes still have boundaries, meaning that some filaments will run out of the FOV.

### Vectorization of actin filament network

Actin filament vectorization from montage tomograms was performed in Amira 2020.2 (Thermo Fisher Scientific) (Rigort et al., 2012; Stalling et al., 2005). Briefly, SOFIMA-stitched cryo-CARE and IsoNet2 denoised montage tomograms were imported into Amira, processed with a local non-means filter (NLM parameters used-Interpretation: XY planes, Spatial standard deviation:3, Intensity standard deviation: 0.2, Search window: 10px, Local neighborhood: 3px), and subjected to filament tracing. For the initial cylinder cross-correlation step, a cylinder length of 360 Å, angular sampling of 5°, a cylinder mask radius of 40 Å, and an outer cylinder radius of 35 Å (corresponding to actin filament radius) was used. Filaments were then traced using the Trace Correlation Lines module with the following parameters: minimum seed correlation 160-180 (cell1 montage#1: 162, cell2 montage#1: 175, cell2 montage#2:170, cell2 monatge#3: 170, and fragment1 montage: 162), minimum continuation quality 100, direction coefficient 0.3, minimum distance 110 Å, minimum length 360 Å, and search-cone minimum step size 10%.

The initial filament tracing obtained from above steps contained false-positive filaments above and below the true actin network, arising from denoising and missing-wedge related artifacts. These filaments were easily identified via inspection of XZ/YZ views and were manually removed in Amira using the Filament Editor module. Residual false positives within the network, primarily corresponding to missing-wedge artifacts (near-Z-aligned traces), were removed using our previously published computational toolbox by excluding filaments oriented within 0-20° of the Z axis (G. Dimchev et al., 2021). Finally, the curated filament vectors were resampled to a fixed spacing of 10 nm to ensure uniform point density along traces using a custom python script. The resulting 10 nm resampled coordinate files (.csv) were used for downstream quantitative actin network analysis.

### Quantitative actin network analysis

Analysis of actin network parameters was performed using custom scripts (see Code availability).

### Actin concentration within the imaged cell volume

The total contour length of all traced actin filaments was obtained by summing the Euclidean distances between consecutive coordinates across every filament in the reconstruction. Total actin content was estimated by dividing the summed filament length by the monomer pitch of 2.7 nm, thereby estimating the number of actin subunits (N_monomers).

To determine the cytoplasmic volume of the imaged lamellipodia regions, two independent geometric estimates were used. An alpha-shape envelope of the filament point cloud provided a concave lower-bound volume estimate, whereas a Gaussian-blurred isosurface of the filament point cloud provided a smoothed upper-bound estimate. The mean of these two values was taken as the best approximation of the effective imaged cell volume.

Actin concentration was calculated as C_actin = N_monomers/ (N_A x V_cell), where N_A is Avogadro’s number and V_cell is the effective cell volume converted to litres. Concentrations were reported in µM.

### Filament length distribution and projected x-y and z components

For every filament, the total contour length was computed as the sum of Euclidean distances between consecutive filament coordinates. The resulting filament traces were then projected onto the x-y plane (Figure 2, S6, S7, and S8 C, D) and onto the z axis (Figure 2, S6, S7, and S8 E, F) to determine the horizontal and vertical components of the length distribution. Histograms of total contour length, x-y-projected length, and z-projected length were generated to visualize spatial anisotropy of filament extension within the cell. **Filament length binning and filament counts**

Filaments were grouped into predefined length intervals, typically using 0.2 µm-wide bins unless otherwise specified. The number of filaments within each bin was counted and converted to relative frequencies to compare the abundance of short, intermediate, and long filaments across datasets or spatial regions. Mean and maximum filament lengths were also reported for each dataset.

### Endpoint classification and barbed, pointed-end distributions

Filament endpoints were operationally classified as barbed or pointed based on their distance to the manually defined leading-edge contour. For each filament, the endpoint closer to the leading edge was assigned as the barbed end, and the opposite endpoint was assigned as the pointed end. Two-dimensional heatmaps were generated from the coordinates of inferred barbed and pointed ends. Relative-frequency histograms were also generated to quantify differences in endpoint distribution between manually specified protrusive and retracting regions of the cell.

### Spatial heatmaps and distance-to-leading-edge profiles

Local filament density, endpoint density, and filament orientation were mapped in two dimensions using spatial binning. Filament density was calculated for each bin from the number of filament segments or coordinate points divided by the bin area. Endpoint density was calculated as the number of endpoints per bin area, and local orientation was calculated from segment vectors within each bin.

For spatial comparisons across the lamellipodium, the distance from each heatmap pixel to the manually traced leading-edge contour was computed. Mean density or orientation values were then plotted as a function of distance from the leading edge to quantify spatial changes in actin organization across the lamellipodium.

### Quantification of Arp2/3 branch junctions and connectivity analysis

#### Mapping Arp2/3 branch junctions within montage tomograms using template matching

For template matching, non-denoised stitched montage tomogram cell2 montage#1 was used. This raw stitched montage tomogram was rendered with the SOFIMA solved mesh obtained from the IsoNet2-denoised stitched volume, while preserving the original raw tomographic signal for template matching. This montage was rotated by 90° about the Z-axis to match the wedge mask applied by PyTom (Hrabe et al., 2012), and then the empty edges of the volume around the stitched montage tomogram were trimmed off. The overall workflow used for Arp2/3 branch-junction identification and refinement is summarized in figure S10.

To generate an initial Arp2/3 search reference, template matching was performed in Dynamo v1.1.33 (Castaño-Díez et al., 2012) on 80 individual montage tiles from the keratocyte fragment using an Arp2/3 branch-junction reference derived from our previously published structure (EMD-11869) (Fäßler, Dimchev, et al., 2020). Approximately 3,800 identified branch junctions were averaged in RELION 4.0 (Scheres, 2012; Zivanov et al., 2022) to generate an initial subtomogram average (STA).

Template matching was then performed using a custom PyTom (version 0.9.0) pipeline (Hrabe et al., 2012). Per-tilt defocus and electron-dose profiles were generated by averaging AreTomo3 (Peck et al., 2026) estimates across montage tiles (median inter-tile standard deviation of 0.16 µm) and approximating accumulated dose from the dose-symmetric tilt scheme, respectively. These per-tilt CTF profiles were supplied as input to PyTom to account for CTF and dose-dependent effects during template matching.

The contrast normalization and 8-bit intensity quantization applied during montage preprocessing and stitching altered the intensity values of the montage tomogram relative to those used to generate the RELION-derived average. This was reflected in broad, poorly localized cross-correlation (CC) peaks at known Arp2/3 branch junctions, reducing the specificity of candidate picking. To address this, we developed a custom Python script to optimize the template-matching parameters using a dataset-specific grid search. Ten clearly identifiable branch junctions lying in the XY plane were selected from the stitched montage and cropped into 128-voxel subvolumes. The script systematically evaluated combinations of template handedness, low-pass and high-pass filtering cutoffs, and masking parameters by performing template matching on each of the ten subvolumes. Both the original and mirrored templates were tested, as the handedness of the stitched volumes could not be assumed. The resulting CC score maps were assessed using the intensity and rank of the peak associated with each known branch junction, relative to other local maxima and the surrounding background. This enabled the identification of parameter combinations that produced strong, well-localized peaks at known branch locations. The selected filtering parameters were a low-pass cutoff of 45 Å and a high-pass cutoff of 123 Å. The selected template handedness and mask were then used for template matching of the full stitched montage volumes.

To manage computational overhead and overcome GPU memory limitations, the stitched montage volume was split internally into overlapping subvolumes during PyTom template matching using the –volume-split option (16×16×2 grid). To prevent false-positive picks on the outer boundaries stitched montage tomogram, a custom script was used to detect the boundaries and generate a soft-edged mask with a 20-pixel taper, which was applied to the cross-correlation (CC) score volume prior to particle extraction.

To overcome the computational bottleneck of performing iterative global maximum searches on the large montage volume, a custom candidate extraction script adapted from PyTom’s pytom_extract_candidates.py was implemented. The script partitioned the score volume into 128³ voxel blocks and tracked the highest-scoring peak from each block using a global priority queue. When the highest overall peak was selected from the queue, a spherical mask with a diameter of 155 Å was applied around the selected peak of the CC score volume to prevent duplicate picking, and the priority queue was updated to identify the next highest-scoring peak. The CC-score cutoff was determined empirically for this dataset in a preliminary picking and classification analysis. The threshold was lowered until the number of candidates classified as branch junctions plateaued, and the resulting cutoff was applied to the full candidate set. The final list of unique candidates was saved to a single STAR file.

Candidates were cropped using a custom Bash script calling Dynamo’s dtcrop command (36^3^-voxel box size; (Castaño-Díez et al., 2012)). The cropped subvolumes were converted to MRC format, contrast-inverted using relion_image_handler, assembled into a RELION-4-compatible STAR file (Scheres, 2012; Zivanov et al., 2022). Stitching and cropping broke the correspondence between particles and their original tilt-series images, so particle-specific CTF information could not be recovered for conventional CTF correction in RELION. A synthetic 3D missing-wedge mask corresponding to the −50° to +50° tilt range was therefore supplied as each particle’s rlnCtfImage to represent the sampled angular range during classification and refinement.

Initial 3D classification was performed with local alignment, followed by duplicate removal using a minimum inter-particle distance of 60 Å to exclude particles that converged to the same location. Subsequently, 3D classification without alignment was alternated with 3D auto-refinement, during which particle orientations and translations were optimized, to separate true branch junctions from actin filaments and false-positive picks. This yielded 6,954 particles. Manual inspection of 250 randomly selected particles showed that 92% were correctly identified branch junctions (Figure S11A).

### Arp2/3 branch junction distribution analysis

To calculate Arp2/3 branch junction distribution with respect to actin filament organization in cell2 montage#1, the two coordinate files (Ap2/3 branch junction positions coordinates and filament coordinates) were first registered to a common reference frame. The non-denoised montage tomogram was rotated and trimmed (as described above), wheras the Amira filament coordinates were obtained from the unrotated IsoNet2-sticthed montage tomogram. Therefore, filament coordinates were transformed to account for the changes in orientation and coordinate origin introduced by rotation and trimming of the tomogram.

To do this, branch junction positions (particles.star file (n = 6,954)) and the transformed filament coordinates of cell2 montage#1 were projected onto the XY plane. The cell outline was defined from the filament coordinates and the filament points were 2D projected and counted on a grid of 50 nm squares, by marking squares containing filament points as cellular region. Small isolated specks were removed, gaps at the rear of the cell due to lower filament density in the projected image were filled, and the largest connected region was kept as the cell outline (IsoNet2 denoising allowed stitching of all the tiles from cell2 montage#1, hence the 2D projected area of the cell outline is 37.7 µm²). We used the filaments rather than the branch junctions to define the cell outline, because the filaments cover the whole montage, whereas junctions become sparse towards the cell interior. To find the leading edge, we first determined the direction in which the cell front points. The 2D projection of branch junction positions was rotated in the image plane in steps of 2° over a half circle. At each angle we counted how many junctions fall into the most crowded band of 200 nm width perpendicular to the viewing axis. Because the junctions are concentrated in a band along the leading edge, branch junction count is highest when the axis is perpendicular to the leading edge. Of the two ends of this axis, the front is the one where the junction density stops abruptly; at the other end the density tails off slowly into the cell interior. The leading edge was then traced along the front side of the cell outline: for each 50 nm wide strip running across the front direction, we took the outermost point of the outline.

For every point inside the cell outline we calculated the shortest distance to the leading edge, and divided the cell into stripes of 500 nm width according to this distance. Branch junctions were assigned to stripes by their distance from the leading edge; junctions outside the cell outline (34 of 6,954, or 0.48% of the total) were not used. Branch junction density was calculated as the number of junctions in a stripe divided by the stripe area. Filament length was calculated as the 3D path length of the traced filaments. Filament density was calculated as the total filament length in a stripe divided by the stripe area, and filament length per branch junction as the total filament length in a stripe divided by the number of junctions in the same stripe. Stripes up to 6 µm from the leading edge are shown. To compare how fast the two densities, change, each was divided by its value in the first 500 nm stripe and plotted against distance from the leading edge.

### Arp2/3 branch junction-based connectivity analysis

Template-matched Arp2/3 branch junctions and Amira-traced actin filaments were then used to identify potential mother–daughter filament connections. Candidate connections were identified geometrically. The mother-filament segment was required to lie within 12 pixels (16.2 nm) of a potential branch point, while the candidate daughter endpoint had to lie within 20 pixels (27.0 nm). The larger distance tolerance for daughter endpoints accommodated filament traces terminating short of the junction, while the subsequent STA-based orientation filter provided an additional constraint on connection assignments. The angle between the local mother- and daughter-filament directions was required to fall within 40–90°, this yielded 2,690 geometrically accepted mother-daughter connections. Not all branch junctions could be assigned to mother–daughter filament pairs. This was potentially due to short daughter-filament stumps (Figure S11B) that were not detected due to being below or close to the minimum tracing length of 36 nm or filaments not being identified during tracing at all.

Candidate mother-daughter connections were then further evaluated against the particle orientations determined during subtomogram averaging (STA). Reference mother- and daughter-filament directions, defined in the local coordinate system of the averaged particle, were transformed into the tomogram coordinate frame using the inverse of the rotation defined by the RELION Euler angles. The predicted directions were compared with the corresponding local mother-filament axis and inward-pointing daughter-filament tangent. Connections with angular discrepancies of ≤25° for both directions were retained. This provided an independent geometric consistency check on the proposed connections. This all-link orientation filter retained 2,236 connections out of the geometrically identified 2,690.

To assess connection accuracy, a random sample of 200 retained mother-daughter connections was inspected manually. Each was assessed for correct assignment of the mother filament, daughter filament and branch point, and for whether the proposed connection represented a genuine filament junction. Manual validation showed that 99% of mother-daughter connections met all these criteria. The retained connections were then represented as directed edges between traced filaments, defining connected components that were treated as filament trees for subsequent analysis. The resulting connected filament trees were visualized in their original spatial positions within the full montage coordinate system. For generation analysis, the root mother filament was assigned generation 0, its directly connected daughters’ generation 1, and subsequent daughter filaments were assigned successively higher generations. The maximum generation depth was calculated for each tree and used to compare and visualize network architecture across the montage.

| REAGENT or RESOURCE | SOURCE | IDENTIFIER |
|---|---|---|
| Chemicals, peptides, and recombinant proteins |  |  |
| DMEM | Gibco | Cat#31966047 |
| START medium | *(Mueller et al. 2017)* |  |
| Fish Ringer | *(Mueller et al. 2017)* |  |
| Steinberg medium | *(Mueller et al. 2017)* |  |
| RUNNING medium | *(Mueller et al. 2017)* |  |
| Cytoskeleton buffer | *(Fäßler et al. 2023)* |  |
| PBS | Gibco | Cat#20012068 |
| PIPES | Sigma-Aldrich | Cat#P8203 |
| Potassium chloride (KCl) | Sigma-Aldrich | Cat#60103 |
| Sodium chloride (NaCl) | Sigma-Aldrich | Cat#S7653 |
| Sodium bicarbonate (NaHCO<sub>3</sub>) | Sigma-Aldrich | Cat#S6297 |
| Calcium chloride dihydrate (CaCl<sub>2</sub>·2H<sub>2</sub>O) | Sigma-Aldrich | Cat#223506 |
| Tris-HCl | Carl Roth | Cat#9090.3 |
| Calcium nitrate tetrahydrate (Ca(NO<sub>3</sub>)<sub>2</sub>·4H<sub>2</sub>O) | Sigma | Cat#C1396 |
| Magnesium sulfate hexahydrate (MgSO<sub>4</sub>·7H<sub>2</sub>O) | Sigma-Aldrich | Cat#M1880 |
| EGTA | Sigma-Aldrich | Cat#E3889 |
| Trypsin-EDTA, 0.25% | Gibco | Cat#25300054 |
| Trypsin inhibitor | Sigma-Aldrich | Cat#T6522 |
| Fibronectin | Sigma-Aldrich | Cat#11051407001 |
| Staurosporine | Sigma-Aldrich | Cat#S4400 |
| Paraformaldehyde | Merck | Cat#P6148 |
| Triton X-100 | Sigma-Aldrich | Cat#X100 |
| Fluoromont-G with DAPI | Invitrogen | Cat#00-4959-52 |
| Glutaraldehyde | Electron Microscopy Sciences | Cat#E16220 |
| Phalloidin | Merck | Cat#P2141 |
| Phalloidin-488 | ATTO-TEC | Cat#AD488-81 |
| Magnesium chloride (MgCl<sub>2</sub>) | Sigma | Cat#M8266 |
| Chicken serum | Sigma-Aldrich | Cat#C5405 |
| Experimental models: Organisms/strains |  |  |
| Zebrafish: wild-type AB/TL Zebrafish (~2 years old) | ISTA Zebrafish facility |  |
| Other |  |  |
| Glass coverslips | Bartelt | Cat#9.161 061 |
| Glass slides | Lactan | Cat#H867.1 |
| 6-well plates | BIOMEDICA | Cat#TP-92696 |
| 12-well plates | BIOMEDICA | Cat#TP-92412 |
| 96-well plates | BIOMEDICA | Cat#TP-92696 |
| Quantifoil SiO<sub>2</sub> R2/2 Ti 200 mesh grids | Quantifoil Micro Tools | Cat#N1-S16nTi20-01 |
| Quantifoil SiO<sub>2</sub> R1.2/20 Au 150 mesh grids | Quantifoil Micro Tools | Cat#N1-S20nAu15-50 |
| 3D-printed grid holders |  | - |

## Supplementary tables

**Table S1:**
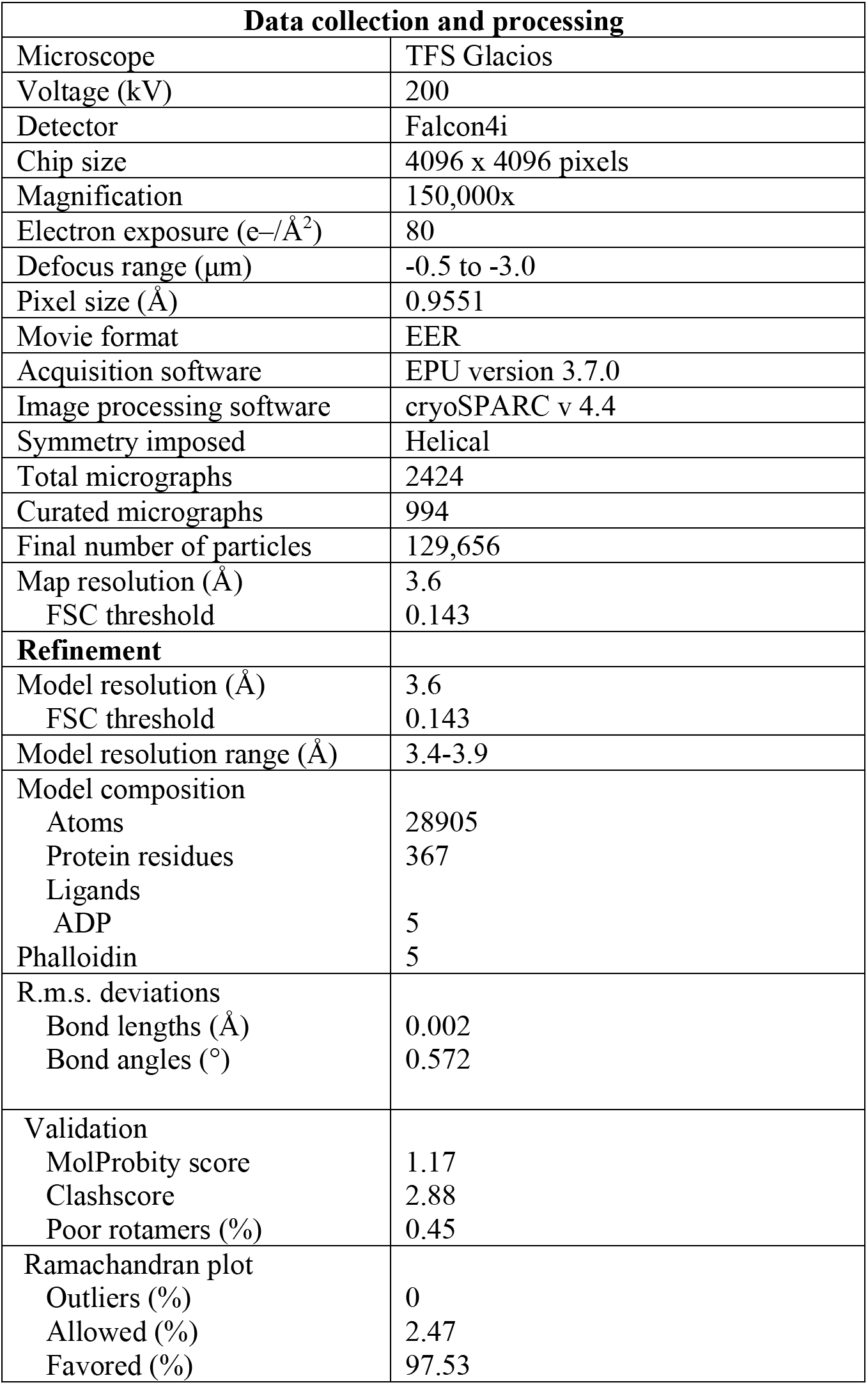
Single-particle data collection, processing and modeling of zebrafish F-actin.

| Data collection and processing |  |
|---|---|
| Microscope | TFS Glacios |
| Voltage (kV) | 200 |
| Detector | Falcon4i |
| Chip size | 4096 x 4096 pixels |
| Magnification | 150,000x |
| Electron exposure (e–/Å<sup>2</sup>) | 80 |
| Defocus range (μm) | -0.5 to -3.0 |
| Pixel size (Å) | 0.9551 |
| Movie format | EER |
| Acquisition software | EPU version 3.7.0 |
| Image processing software | cryoSPARC v 4.4 |
| Symmetry imposed | Helical |
| Total micrographs | 2424 |
| Curated micrographs | 994 |
| Final number of particles | 129,656 |
| Map resolution (Å) | 3.6 |
| FSC threshold | 0.143 |
| **Refinement** |  |
| Model resolution (Å) | 3.6 |
| FSC threshold | 0.143 |
| Model resolution range (Å) | 3.4-3.9 |
| Model composition |  |
| Atoms | 28905 |
| Protein residues | 367 |
| Ligands |  |
| ADP | 5 |
| Phalloidin | 5 |
| R.m.s. deviations |  |
| Bond lengths (Å) | 0.002 |
| Bond angles (°) | 0.572 |
| Validation |  |
| MolProbity score | 1.17 |
| Clashscore | 2.88 |
| Poor rotamers (%) | 0.45 |
| Ramachandran plot |  |
| Outliers (%) | 0 |
| Allowed (%) | 2.47 |
| Favored (%) | 97.53 |

**Table S2:** Data acquisition parameters used for montage cryo-ET datasets.

| Microscope | FEI Titan Krios |  |  |
|---|---|---|---|
| Voltage [kV] | 300 |  |  |
| Detector | Gatan BioQuantum K3 |  |  |
| Original rectangular chip size | 5760 x 4092 pixels |  |  |
| Cropped square size | 4092 x 4092 pixels |  |  |
| Beam diameter | 1.96 µm |  |  |
| Detector field of view | 1.38 x 1.38 µm |  |  |
| Nominal magnification | 26,000x |  |  |
| Energy-filter | Yes (slit width 20eV) |  |  |
| Acquisition software | SerialEM 4.1.10<br>PACE-tomo 1.4.4 |  |  |
| Defocus [µm] | −6.0 |  |  |
| Acquisition scheme | Dose-symmetric |  |  |
| Tilt range [°] | -50 to +50 |  |  |
| Tilt increment [°] | 2 |  |  |
| Number of frames | 8 |  |  |
| Data set | **Cell1 (1 montage)** | **Cell2 (3 montages)** | **Fragment1** |
| Tile overlap in X and Y | 15% | 10% | 10 % |
| Pixel size [Å/pix] | 3.371 | 3.371 | 3.381 |
| Total dose [electrons/Å²] on one tile | 100 | 100 | 90 |
| Total dose [electrons/Å²] on 4x overlap region | 400 | 400 | 360 |

**Table S3:** Quantitative parameters of actin networks obtained from montage tomograms.

| Cell/montage | Cell1 montage#1 | Cell2 montage#1 | Cell2 montage#2 | Cell2 montage#3 |
|---|---|---|---|---|
| Montage area | 81.6 µm<sup>2</sup> | 37.4 µm<sup>2</sup> | 51.6 µm<sup>2</sup> | 55.4 µm<sup>2</sup> |
| Total number of filaments | 53,632 | 22,468 | 40,216 | 39,544 |
| Cumulative filament length | 1.08 cm | 0.55 cm | 0.87 cm | 0.93 cm |
| Estimated actin subunits | 3,983,723 | 2,041,412 | 3,239,038 | 3,449,019 |
| Estimated network volume isosurface | 7.51 µm³ | 3.98 µm³ | 6.79 µm³ | 6.70 µm³ |
| Estimated network volume alpha shape | 5.27 µm³ | 3.07 µm³ | 5.23 µm³ | 5.43 µm³ |
| Mean F-actin concentration | 1035.59 µM | 960.59 µM | 894.93 µM | 944.14 µM |
| Mean filament length | 0.200 µm | 0.245 µm | 0.217 µm | 0.235 µm |
| Maximum filament length | 4.75 µm | 4.65 µm | 4.78 µm | 4.83 µm |

## Supplementary figures

**Figure S1:**
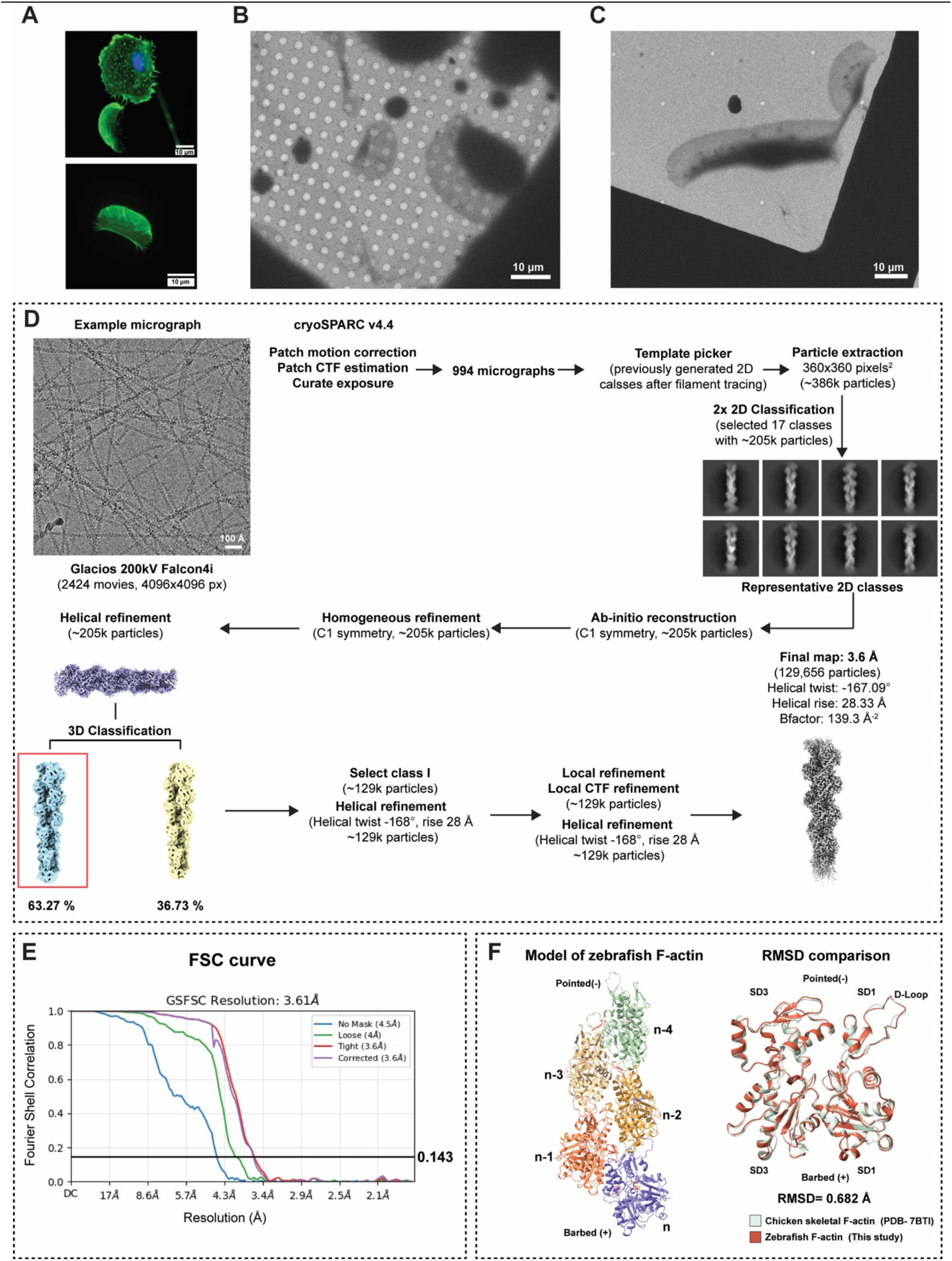
Sample preparation and single-particle cryo-EM validation of keratocyte F-actin. **(A)** Representative fluorescence microscopy images of keratocytes and cytoplasmic fragments on fibronectin-coated glass coverslips after staurosporine treatment. Top: keratocyte forming a cytoplasmic fragment, with an actin-rich fragment separating from the nucleated cell body. Bottom: independently spread cytoplasmic fragment with a broad, polarized, actin-rich lamellipodial region. F-actin was stained with fluorescent phalloidin (green) and nuclei with DAPI (blue). **(B, C)** Representative low-magnification cryo-EM images of vitrified keratocytes and cytoplasmic fragments on fibronectin-coated SiO₂ grids showing thin, spread cellular regions suitable for cryo-ET on a holey SiO₂ R2/2 grid (B), and on a holey SiO₂ R1.2/20 grid (C). (**D**) Single-particle cryo-EM processing workflow for keratocyte actin filaments showing a representative micrograph, 2D and 3D classes, and the final map (see Methods for details). This yielded a final reconstruction at 3.6 Å resolution from 129,656 particles, with a helical twist of −167.09° and a rise of 28.33 Å. (**E**) Gold-standard Fourier shell correlation (GSFSC) curve of the final reconstruction, showing a resolution of 3.61 Å at the 0.143 criterion. (**F**) Left: atomic model of zebrafish F-actin built into the final map, showing five consecutive subunits (n to n−4) and the barbed (+) and pointed (−) ends. An AlphaFold2 model of zebrafish β-actin (UniProt A8WG05) was used as the starting model. Residues 6–372, together with ADP, Mg²⁺, and phalloidin, were fitted into the density using ISOLDE (Croll, 2018) in ChimeraX (Meng et al., 2023; Pettersen et al., 2021) and refined in Phenix (Afonine et al., 2018). Right: superposition of a zebrafish actin subunit (this study, orange) with chicken skeletal F-actin (PDB 7BTI, grey (Kumari et al., 2020)), showing close agreement (RMSD 0.682 Å). Subdomains SD1 and SD3 and the D-loop are labeled. Scale bars: 10 µm (A–C); 100 Å (example micrograph in D).

**Figure S2:**
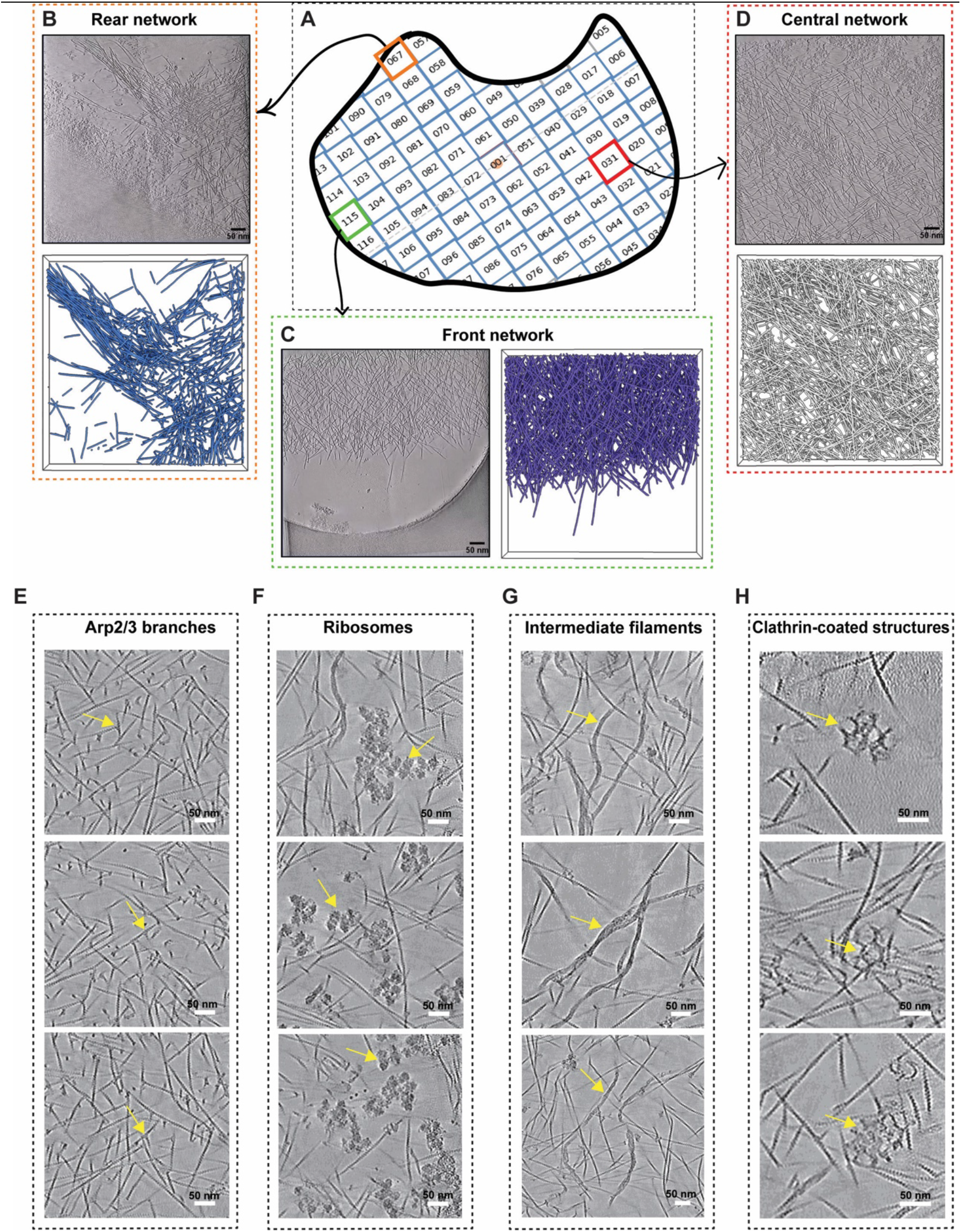
Gallery of different actin architectures and additional molecular entities. **(A)** Schematic of the PACE-tomo montage acquisition on keratocyte fragment#1, showing the numbered tile positions. Colored boxes mark the tiles shown in B (orange, tile 067), C (green, tile 115), and D (red, tile 031). **(B–D)** Representative tiles from different regions of the fragment. Each shows a 1.352-nm slice through the cryo-CARE denoised tomogram, alongside the corresponding vectorized actin filaments rendered in ChimeraX (Meng et al., 2023; Pettersen et al., 2021). **(B)** Network at the rear of the fragment, showing actin bundles; slice (top) and vectorized filaments (bottom). **(C)** Front network at the leading edge; slice (left) and vectorized filaments (right). **(D)** Central network; slice (top) and vectorized filaments (bottom). **(E–H)** Molecular entities resolved in the montage data, shown in 1.352-nm slices through cryo-CARE denoised tomograms: **(E)** Arp2/3 branch junctions, **(F)** ribosomes, **(G)** intermediate filaments, and **(H)** clathrin-coated structures. Yellow arrows mark examples of each. Scale bars: 50 nm.

**Figure S3:**
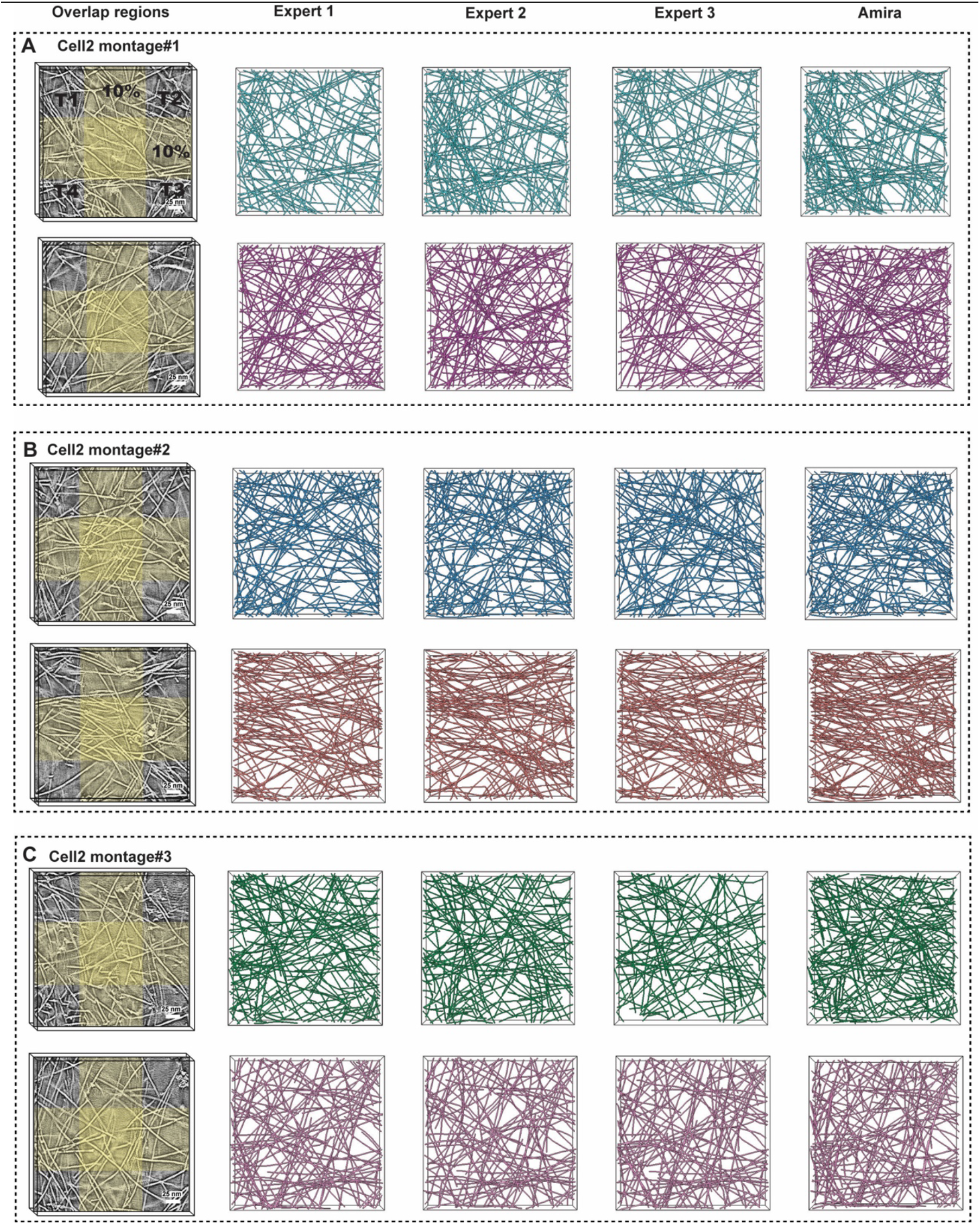
Validation of 3D stitching using manual and semi-automated filament tracing. Comparison of manual and automated actin filament tracing in cropped sub-volumes (400×400 px) from four-tile overlap regions of SOFIMA-stitched cryo-CARE montage tomograms. Sub-volumes were selected from overlap regions that extended beyond the nominal 10% tile-overlap (∼100 px), allowing filaments to be followed from one tomogram into adjacent tomograms. (A-C) Two representative slices at different positions in the montage (thickness 1.35 nm) through overlap-region sub-volumes are shown for each of three independently stitched montages from Cell2: montage#1 (A), montage#2 (B), and montage#3 (C), respectively. In the first example, the four contributing tomograms are indicated as T1-T4, and the 10% overlap region is highlighted in yellow. For each sub-volume, actin filaments were manually traced in IMOD (Kremer et al., 1996), independently by three expert users and by the semi-automated filament-tracing workflow in Amira (Rigort et al., 2012; Stalling et al., 2005). Manual traces from the three experts showed similar filament organization across tile boundaries, and the semi-automated Amira traces reproduced the same overall filament network, although with denser filament detection in some regions. The agreement between manual and automated traces demonstrates that SOFIMA stitching preserved filament continuity across overlap regions and that the stitched volumes were suitable for downstream automated actin filament vectorization. Please note that the expert traces were not done to detect 100% of the filaments but to validate filament continuity across tile boundaries. Scale bars, 25 nm.

**Figure S4:**
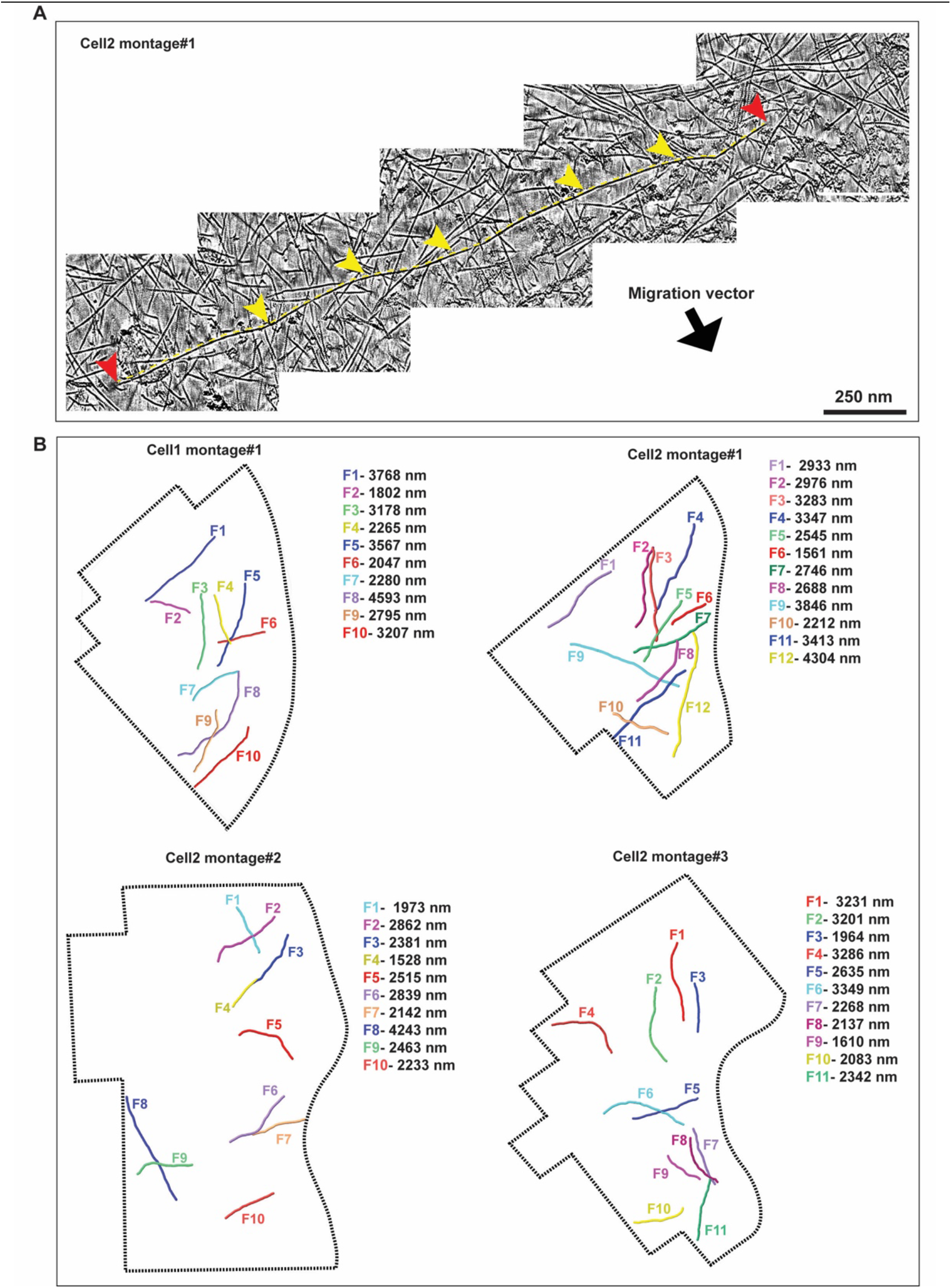
Manual tracing of long actin filaments across multiple tiles in stitched montage tomograms. **(A)** Representative example of a long filament traced through several adjacent tiles in Cell2 montage#1. The filament path is indicated by yellow arrowheads, and the filament ends are marked by red arrowheads. Black arrow indicates the direction of migration of the cell. **(B)** Examples of long filaments manually traced from the stitched montage datasets from Cell1 and Cell2. Individual filaments are shown in different colours, and their measured contour lengths are indicated next to each montage region (black dotted lines indicate montage regions and montage outlines are not for scale). Manual tracing to identify these long filaments was performed in IMOD (Kremer et al., 1996). Scale bars: 250 nm (A).

**Figure S5:**
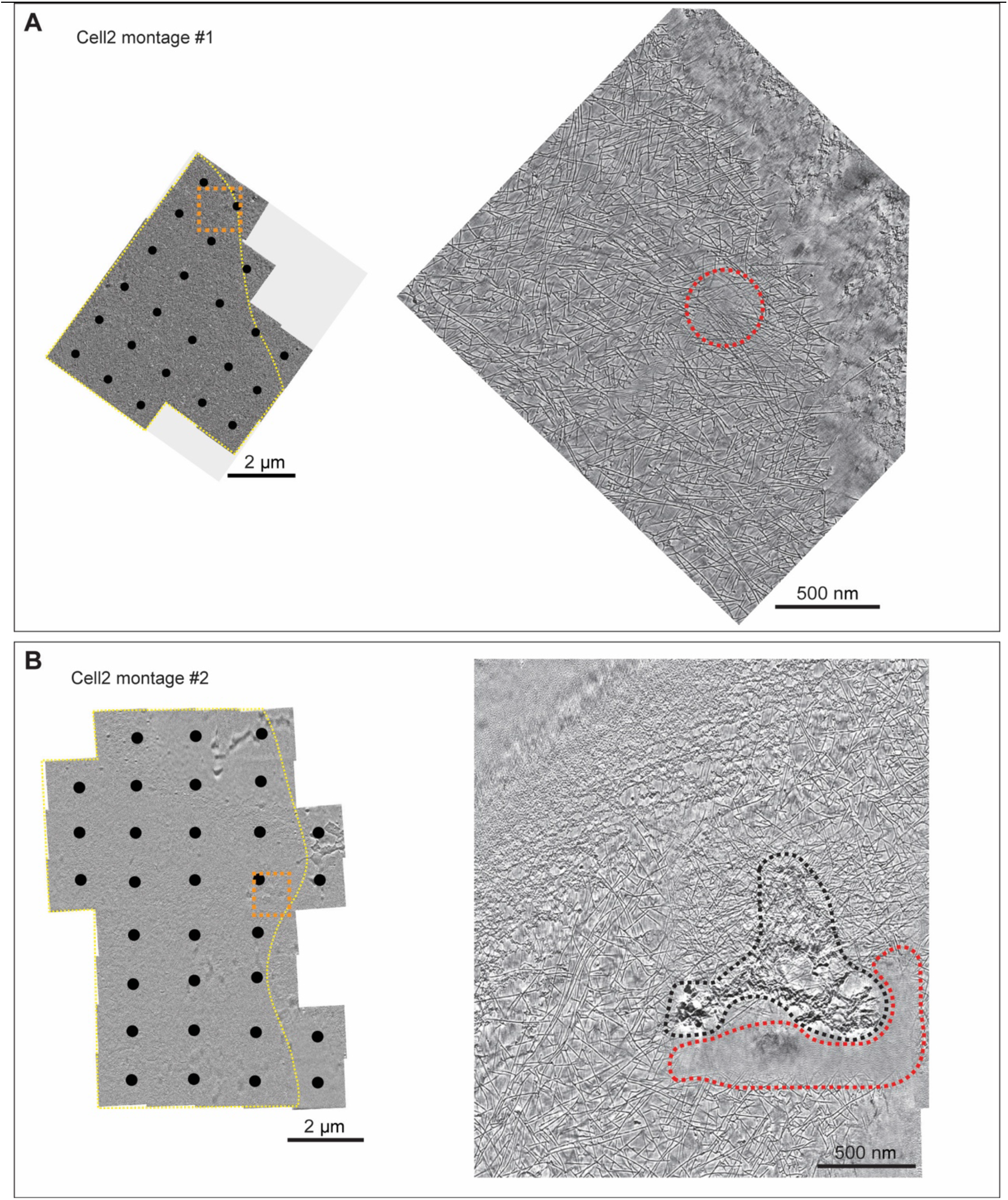
Representative regions of montage tomograms with lower stitching quality. (A) Example region from cell2 montage#1. Left: 1.352-nm slice through the cryo-CARE denoised, stitched montage tomogram. The orange dotted square marks a region of inaccurate stitching. Right: enlarged view of the boxed region, where the red dotted circle marks a tile-overlap region with low stitching quality. (B) Cell2 montage#2. Left: 1.352-nm slice through the cryo-CARE denoised, stitched montage volume, with the orange dotted square marking a region of inaccurate stitching. Right: enlarged view of the boxed region, where the red dotted outline marks a tile-overlap region with low stitching quality caused by adjacent ice contamination (black dotted outline). Black dots mark PACE-tomo acquisition points. Scale bars: 2 µm (left), 500 nm (right).

**Figure S6:**
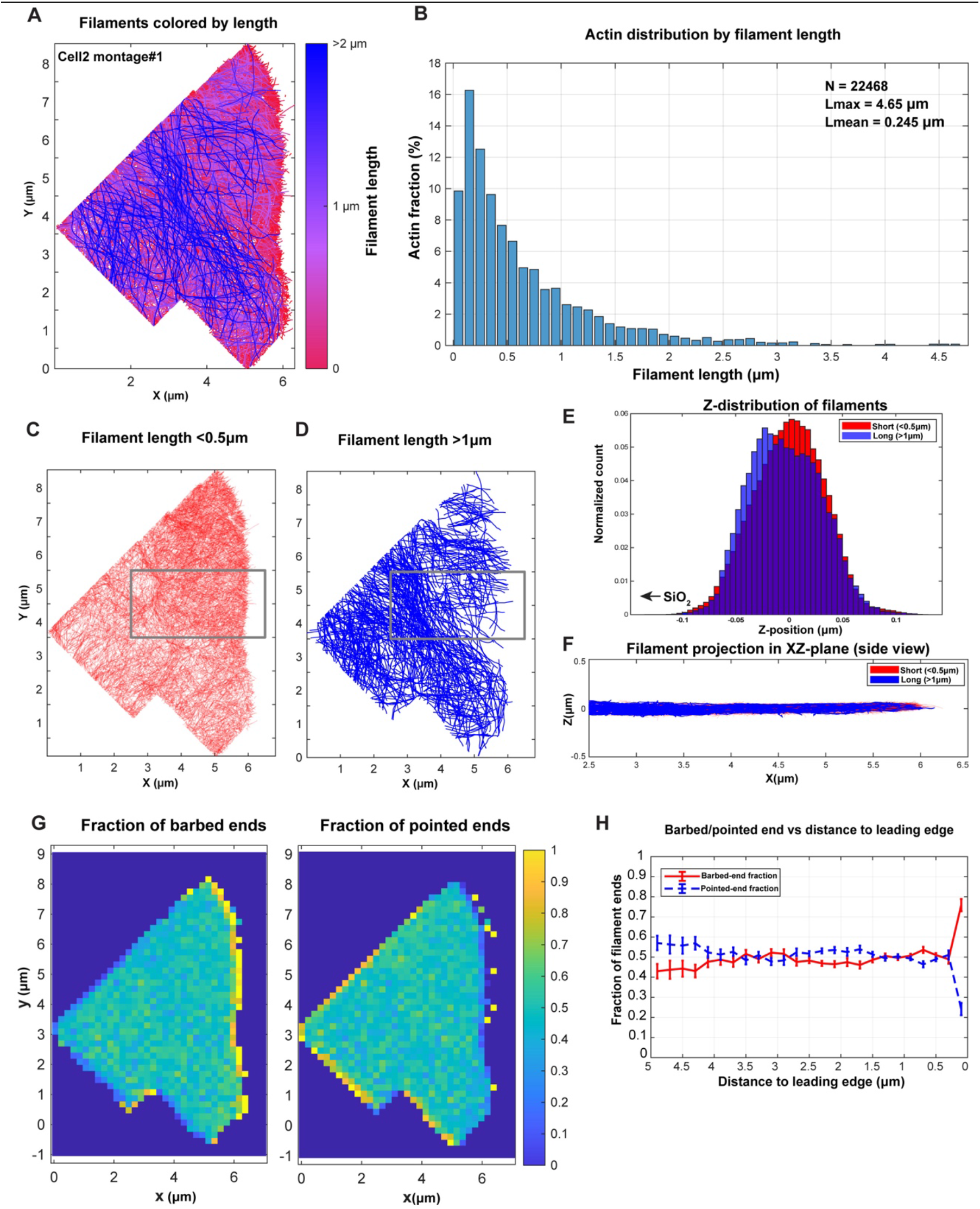
Quantitative analysis of actin filament organization in Cell2 montage#1. **(A)** XY projection of all actin filaments traced in the cryo-CARE-denoised Cell2 montage#1, color-coded according to filament length. **(B)** Actin distribution as a function of filament length, showing the fraction of total actin length in each length bin. The total number of traced filaments (N), maximum filament length (Lmax), and mean filament length are indicated in the plot (Lmean). **(C, D)** XY spatial distribution of filament-length subpopulations. Short filaments, defined as filaments shorter than 0.5 µm, are shown in C, whereas long filaments, defined as filaments longer than 1 µm, are shown in D. **(E)** Z-position distribution of short filaments <0.5 µm and long filaments >1 µm (position of the SiO2 substrate is indicated for orientation). **(F)** XZ projection of short and very long filaments from the boxed regions indicated in C and D. **(G)** Spatial heat maps showing the local fraction of barbed ends and pointed ends across the montage. **(H)** Quantification of barbed-end and pointed-end fractions as a function of distance from the leading edge.

**Figure S7:**
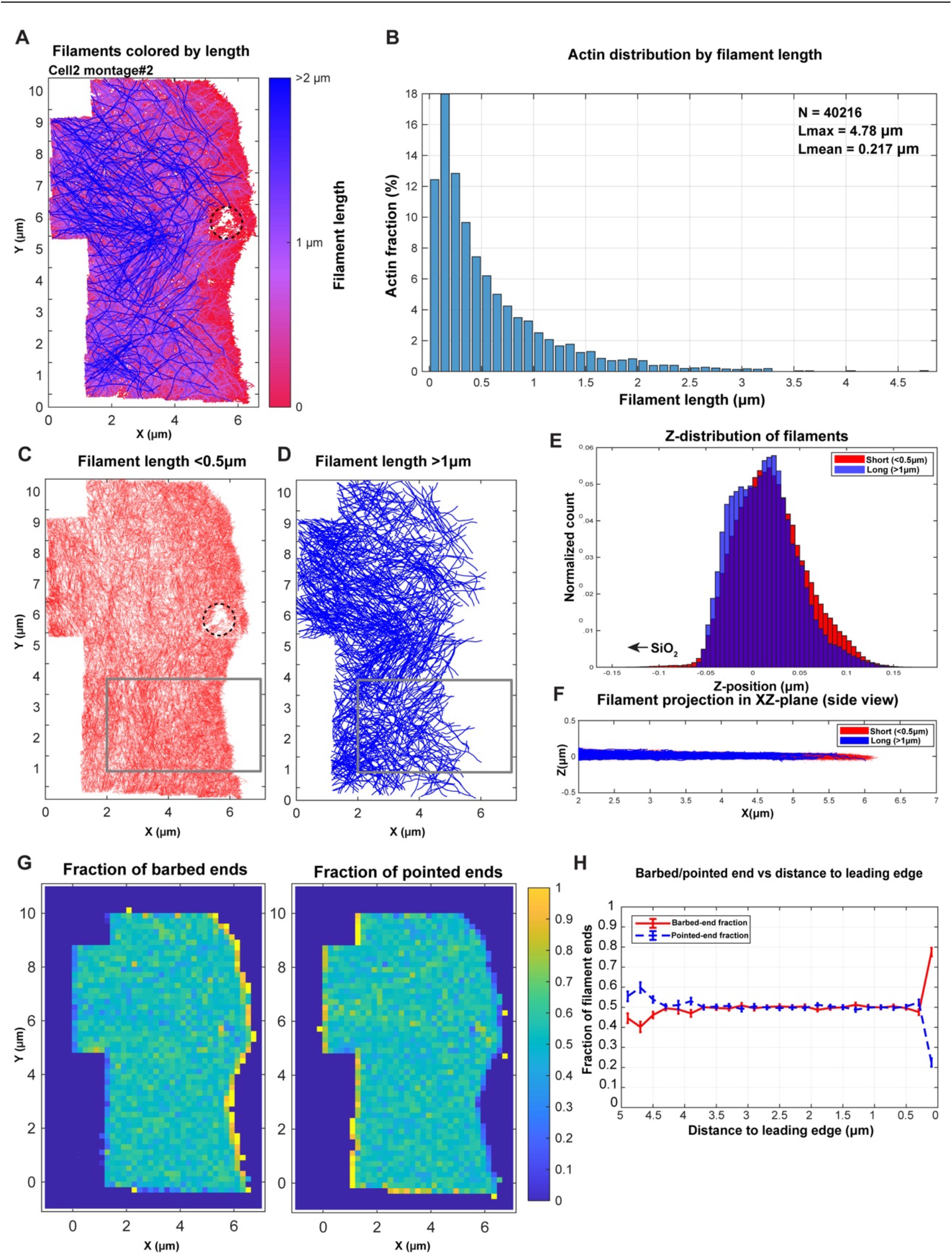
Quantitative analysis of actin filament organization in Cell2 montage#2. **(A)** XY projection of all actin filaments traced in the cryo-CARE-denoised Cell2 montage#2, color-coded according to filament length. A region where filaments could not be traced properly due to ice contamination is marked by a black dashed circle (also in C). **(B)** Distribution of actin as a function of filament length, expressed as the fraction of total actin length in each length bin. The total number of traced filaments (N), maximum filament length (Lmax), and mean filament length are indicated in the plot (Lmean). **(C, D)** XY spatial distribution of short and long filament subpopulations. Short filaments <0.5 µm are shown in C, and long filaments >1 µm are shown in D. **(E)** Z-position distribution of short filaments <0.5 µm and long filaments >1 µm (position of the SiO2 substrate is indicated for orientation). (F) XZ projection of the boxed regions marked in C and D. **(G)** Spatial heat maps showing the local fractions of barbed and pointed ends across the montage. **(H)** Barbed-end and pointed-end fractions plotted as a function of distance from the leading edge.

**Figure S8:**
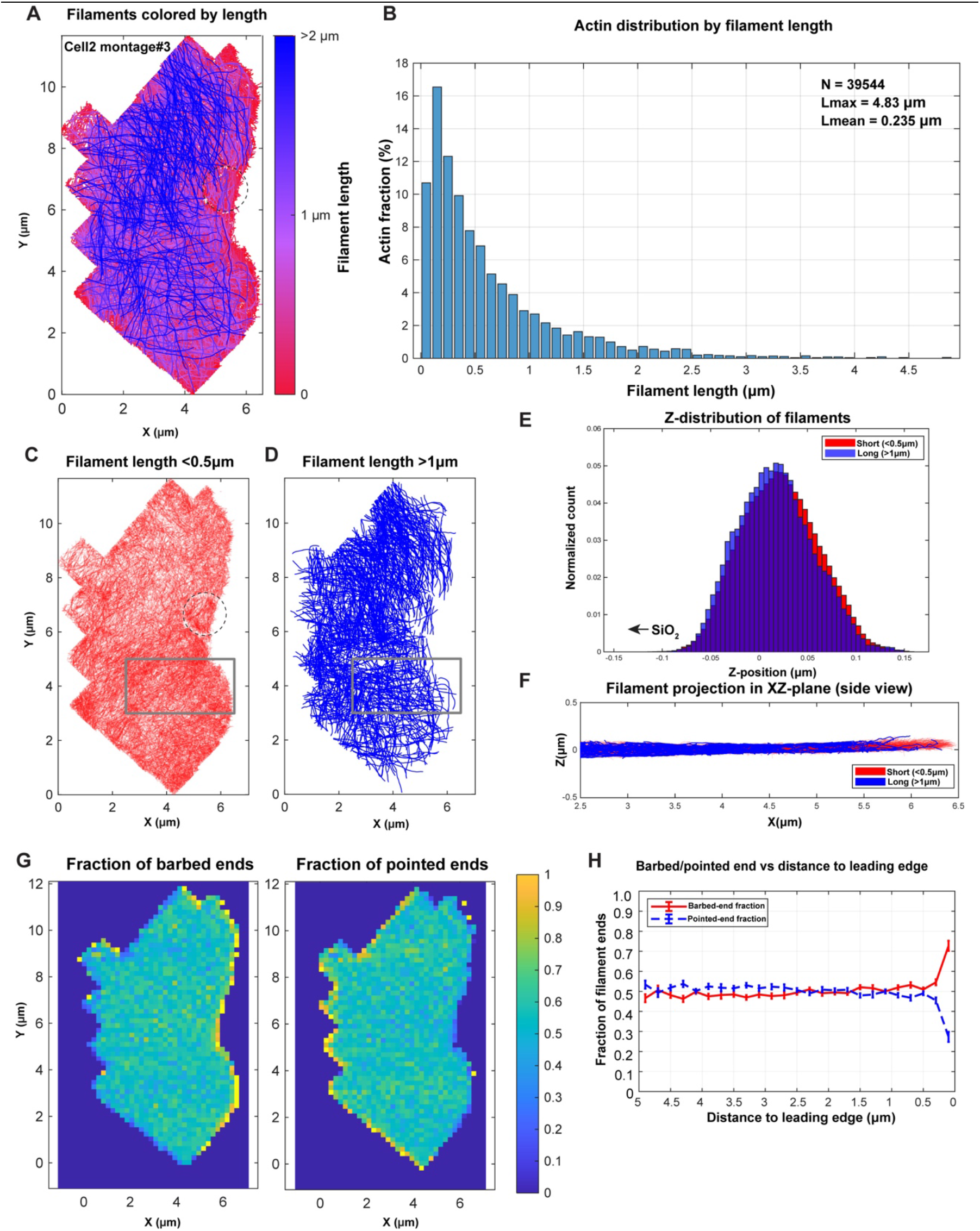
Quantitative analysis of actin filament organization in Cell2 montage#3. **(A)** XY projection of all actin filaments traced in the cryo-CARE-denoised Cell2 montage#3, color-coded according to filament length. The hole edge in the SiO2 grid substrate is marked by a black dashed circle (also in C). **(B)** Actin distribution by filament length, showing the fraction of total actin in each length bin. The total number of traced filaments (N), maximum filament length (Lmax), and mean filament length are indicated in the plot (Lmean). **(C, D)** XY distribution of short and long filament populations. Filaments shorter than 0.5 µm are shown in C, and filaments longer than 1 µm are shown in D. **(E)** Z-position distribution of short filaments <0.5 µm and long filaments >1 µm (position of the SiO2 substrate is indicated for orientation). **(F)** XZ projection of filaments from the boxed regions shown in C and D, comparing the vertical distribution of short and long filaments. **(G)** Spatial heat maps showing the local fractions of barbed and pointed filament ends. **(H)** Barbed-end and pointed-end fractions plotted as a function of distance from the leading edge.

**Figure S9:**
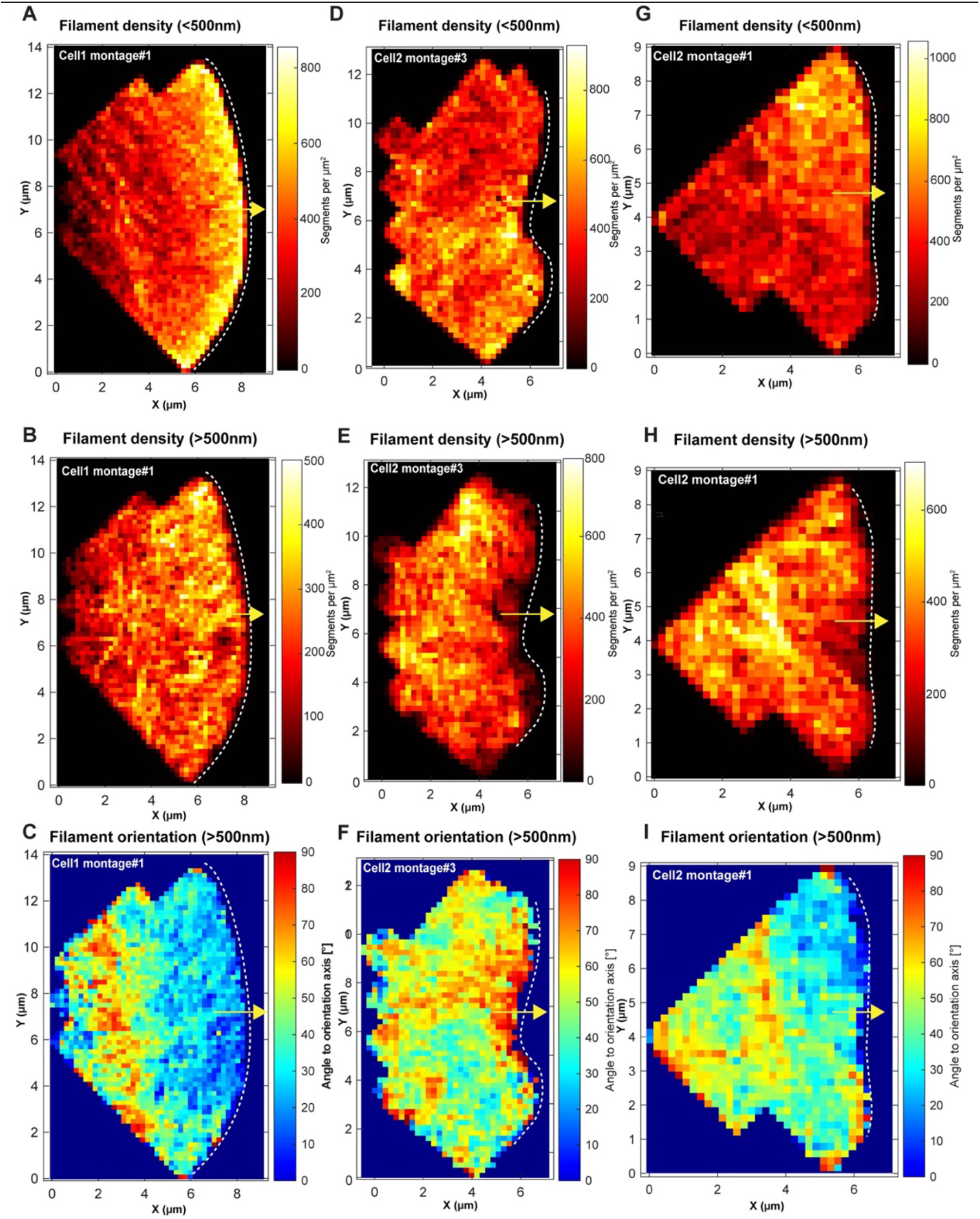
Heterogeneity in filament density and orientation distribution of long and short filaments. **(A, D, G)** Spatial maps of short actin filaments (<500 nm) in three montages from two cells. In all the panels, the yellow arrow indicates the migration direction and the white dashed line indicates the leading-edge shape. Filament density and orientation were mapped in 2D using spatial binning. Density was calculated for each spatial bin (0.2 µm X 0.2 µm) as the number of filament segments divided by the bin area (segments per µm^2^)). Local orientation was calculated from the filament 3D end-to-end vectors within each bin, as the angle relative to the migration direction (yellow arrow). **(B, E, H)** Corresponding spatial maps of long actin filaments (>500 nm) for the montages shown in A, D, G. **(C, F, I)** Filament-orientation distribution of long actin filaments (<500 nm) for the same montages shown in A, D, G. The regions with a convex leading edge show a higher density of short filaments (D, G), consistent with a protrusive-like region. In the neighboring region with a concave or flat leading edge, long filaments are oriented parallel to the leading edge (F, I), consistent with a pausing-like region.

**Figure S10:**
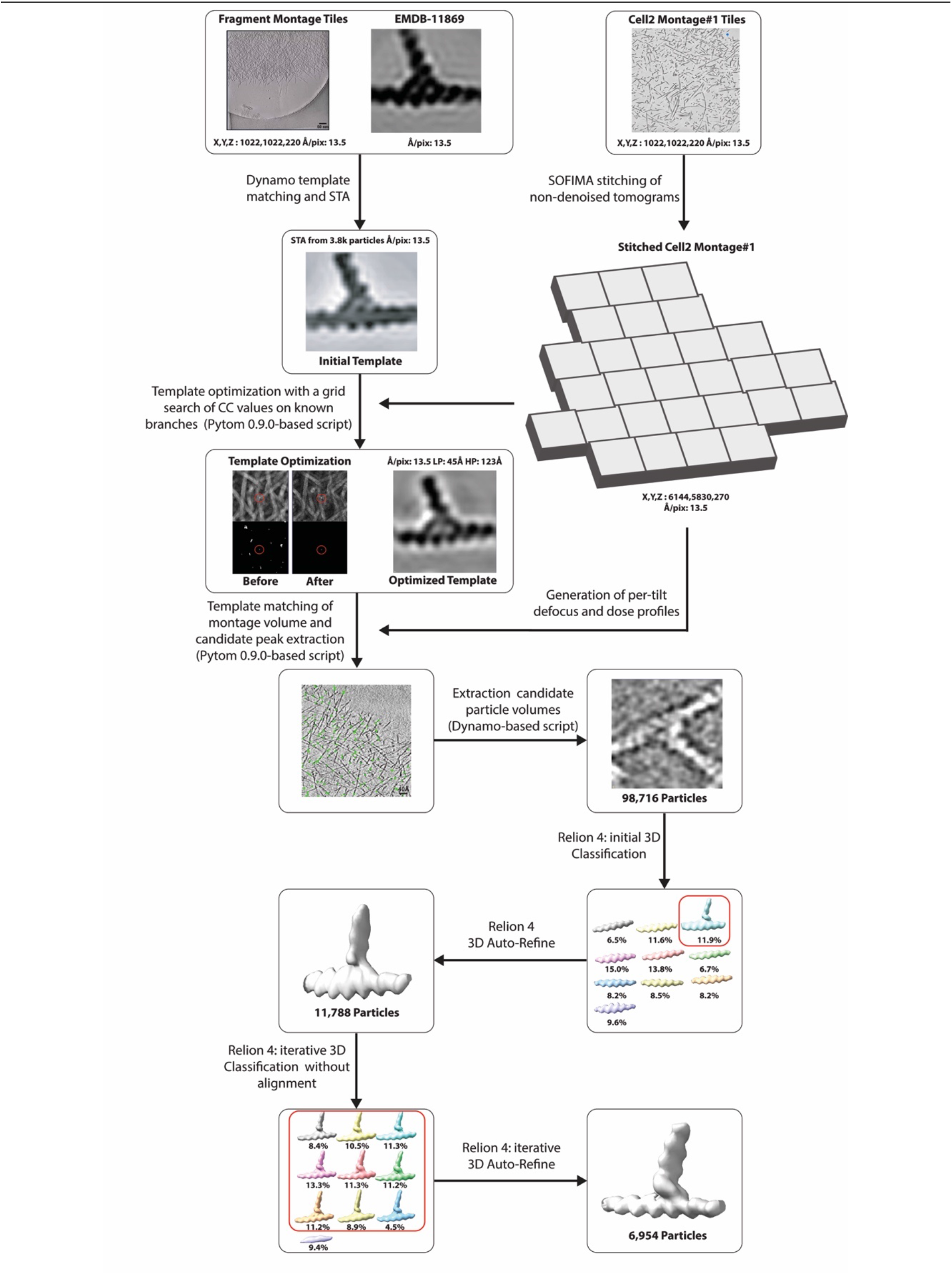
Workflow to identify Arp2/3 branch junctions in the stitched montage tomogram. An Arp2/3 branch-junction structure from our previous work (EMDB-11869) was used for initial Dynamo template matching on 80 individual tomogram tiles from a keratocyte fragment. Approximately 3,800 identified branch junctions were averaged in RELION 4.0 to generate an initial reference. This reference was subsequently optimized using a grid search on ten known branch junctions from the stitched Cell2 montage#1, producing the optimized template used for template matching of the full non-denoised montage with pytom-match-pick. Candidate coordinates were extracted into a STAR file using a custom PyTom-based script, followed by subtomogram extraction and contrast inversion using a Dynamo-based script. Initial 3D classification with local alignment was followed by iterative 3D auto-refinement and classification without alignment to enrich the particle set for true branch junctions. Representative classification results are shown at different stages of iterative 3D classification without alignment and 3D auto-refinement. Red outlines indicate classes selected for subsequent processing, and percentages indicate the particle distribution among classes at each classification stage. The final refinement contained 6,954 branch junctions.

**Figure S11:**
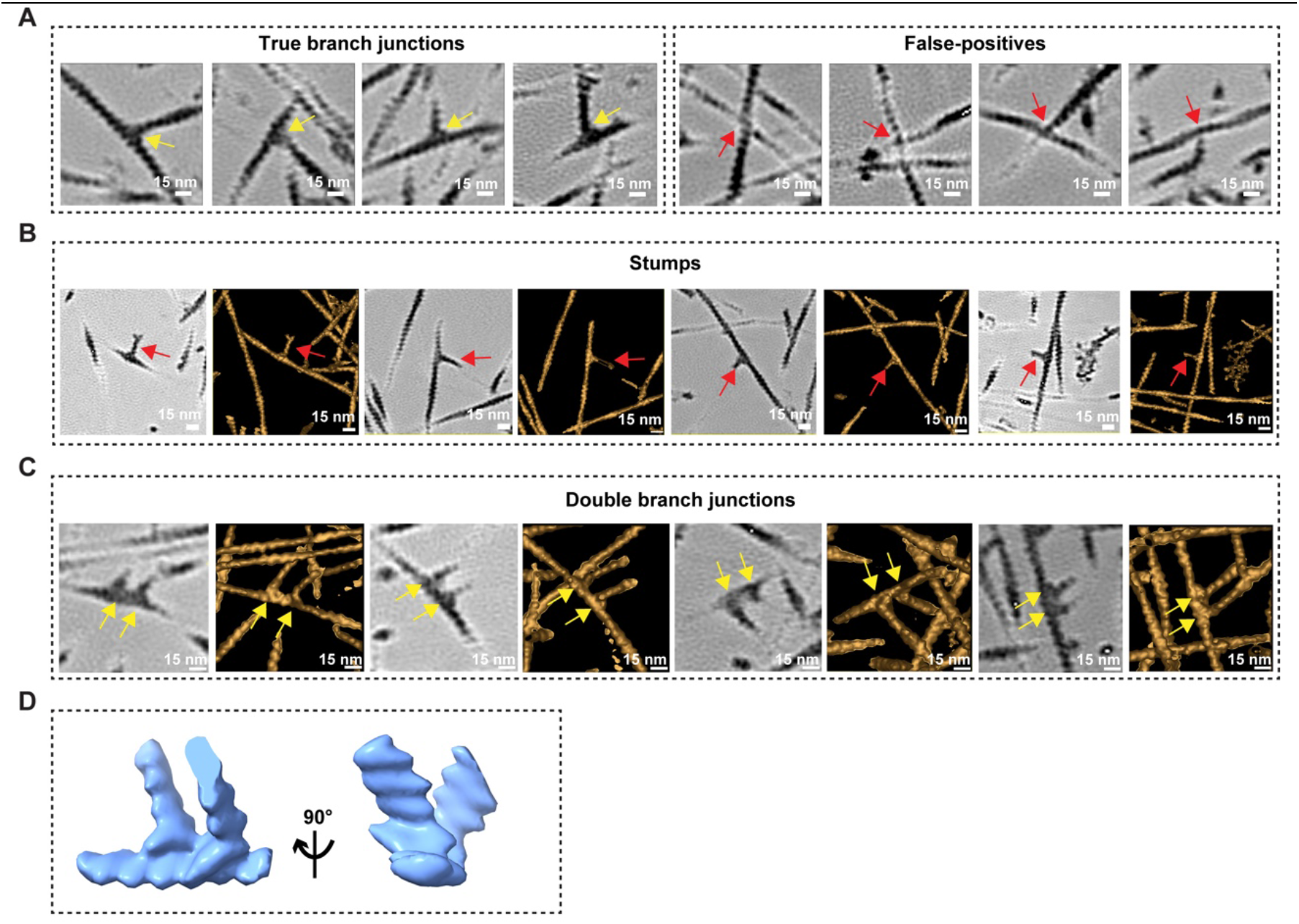
Validation of Arp2/3 branch junction detection and examples of stumps and double branch junctions. Arp2/3 branch junctions were detected by template matching on the misalignment-corrected, raw stitched montage tomogram of Cell2 montage#1. All images shown were cropped at the detected positions from the corresponding IsoNet2-denoised stitched montage tomogram and generated in IMOD (Kremer et al., 1996). **(A)** Representative true branch junctions (left, yellow arrows) and false positives (right, red arrows), shown as 1.352-nm slices. Visual inspection of 250 subtomograms cropped at template-matched positions showed that 92% were true branch junctions and 8% were false positives, mainly filament crossings resembling branch junctions. **(B)** Representative stumps, branch junctions with a very short daughter filament (red arrows). **(C)** Representative double branch junctions, in which two daughter filaments branch from adjacent positions on the same mother filament (yellow arrows). In B and C, each example is shown as a 1.352-nm slice (left) and the corresponding isosurface rendering of 20 slices in Z (27.04 nm; right). **(D)** Isosurface rendering of the sub-tomogram average of the double branch junction from Cell2 montage#1, shown in two views rotated by 90° (Images prepared using ChimeraX (Meng et al., 2023; Pettersen et al., 2021). Scale bars: 15 nm.

**Figure S12:**
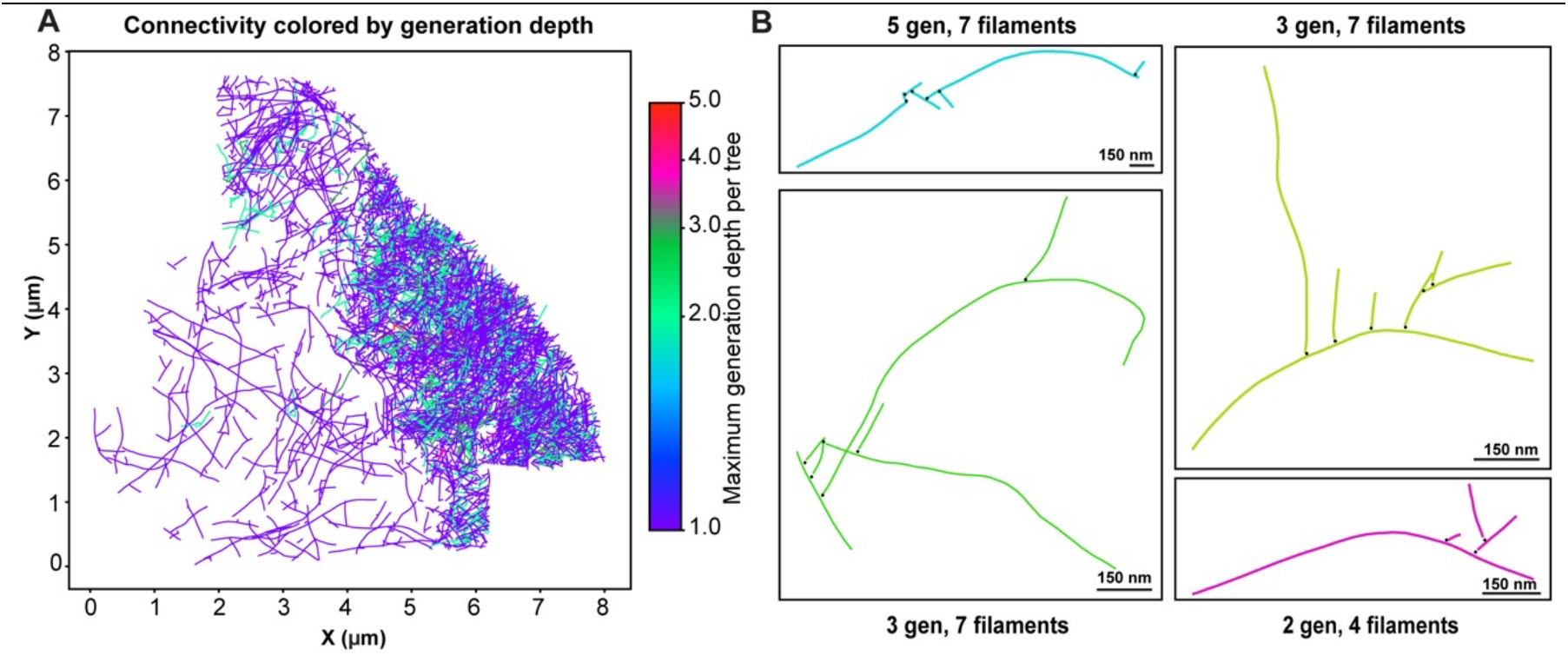
Branch junction-based connectivity depth and topology of filament trees. Actin network connectivity analysis based on Arp2/3 branch junctions and filament coordinates from Cell2 montage#1. **(A)** 2D projection of filament trees, in which filaments connected through Arp2/3 branch junctions are colored by the maximum number of daughter-filament generations per tree (1–5), as indicated by the color bar. **(B)** Selected filament trees illustrating different generation depths and topologies, with each tree shown in a different color. Lines represent filaments and black dots mark the Arp2/3 branch junctions connecting them. The generation depth and number of filaments are indicated for each tree. Scale bars: 150 nm.

**Figure S13:**
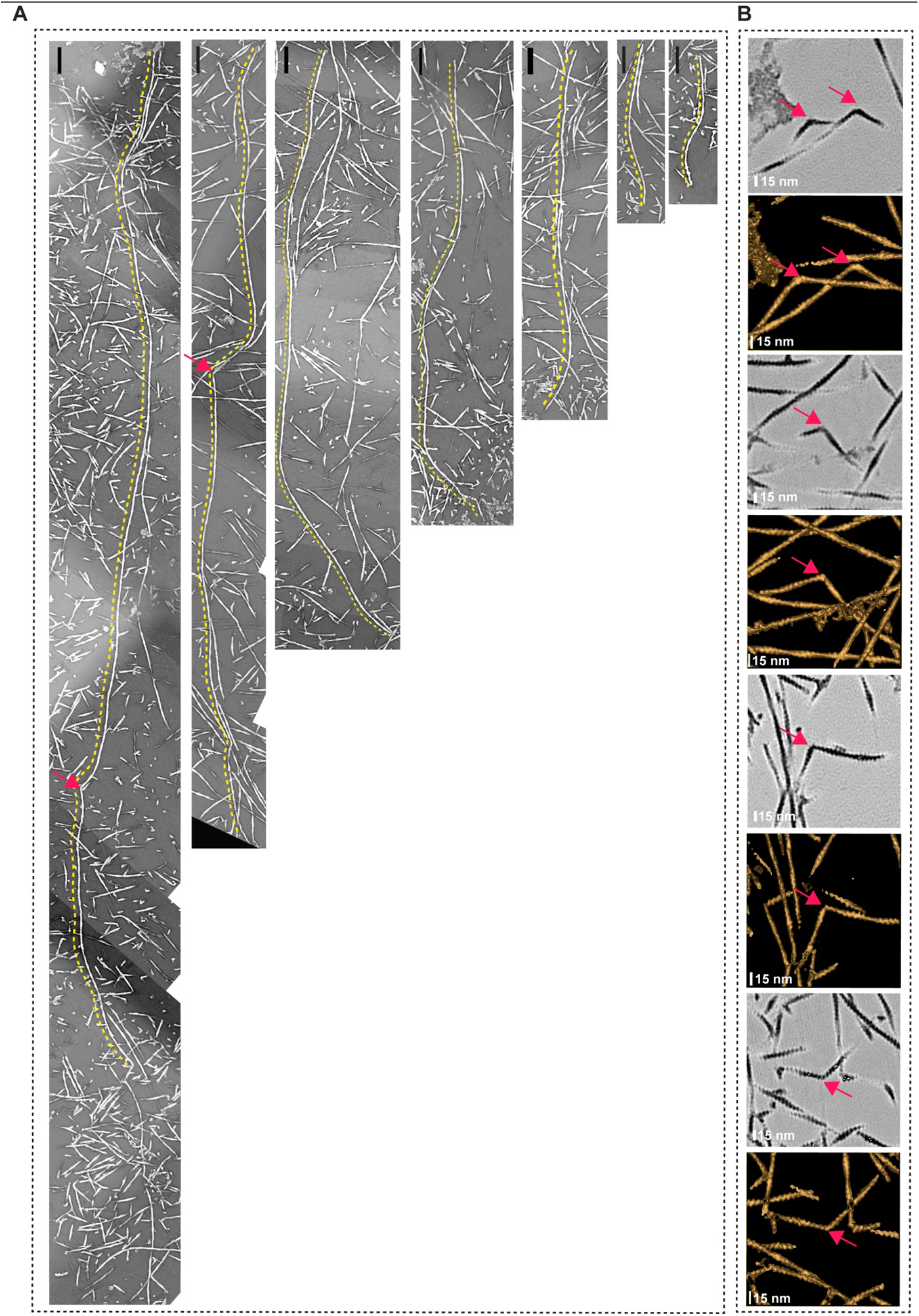
Manually traced long unbranched filaments and filament kinks. **(A)** Seven representative long unbranched actin filaments manually traced in the IsoNet2-denoised, stitched montage tomogram of Cell2 montage#1. Yellow dashed lines mark the traced filaments. Filament lengths from left to right: ∼3.8 µm, ∼3.2 µm, ∼2.3 µm, ∼1.8 µm, ∼1.2 µm, ∼0.6 µm, and 0.5 µm, respectively. **(B)** Representative filament kinks in the IsoNet2-denoised, stitched montage tomogram, shown as four pairs. In each pair, the upper image is a cropped 1.352-nm slice and the lower image is the corresponding isosurface rendering. Isosurfaces include 20 slices in Z (27.04 nm) to show the kinks clearly while excluding overlapping filaments above and below. Pink arrows mark the kinks. All images were generated in IMOD (Kremer et al., 1996). Scale bars: 100 nm (A), 15 nm (B).

**Figure S14:**
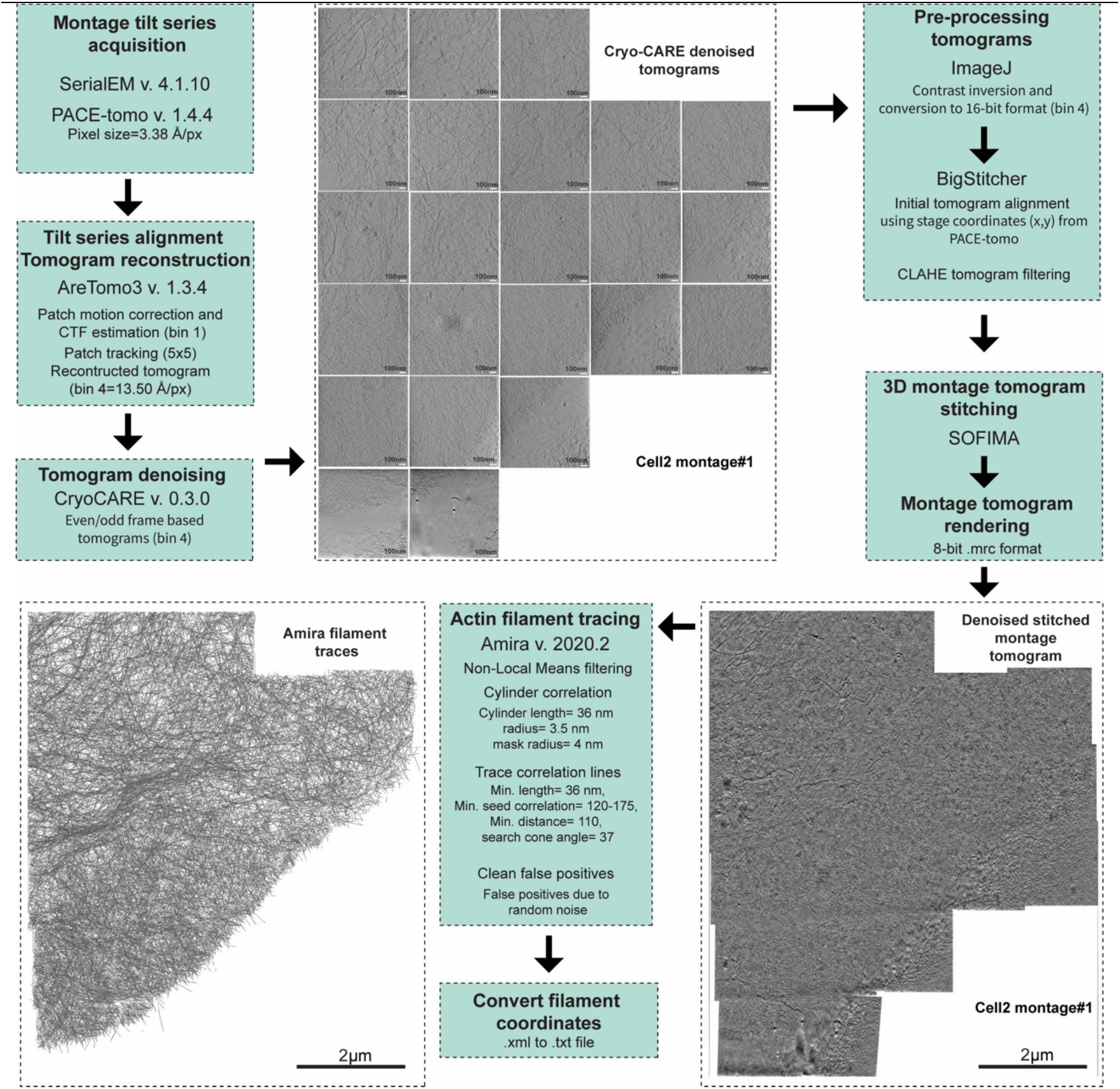
A graphical summary of the workflow for large-scale montage cryo-ET of actin networks. Montage tilt series are acquired in SerialEM and each tile is independently aligned and reconstructed in AreTomo3. Reconstructed tomograms are denoised using Cryo-CARE and then pre-processed in ImageJ by contrast inversion and conversion to 16-bit format. The tile positions from PACE-tomo-based data acquisition are used for initial 3D alignment in BigStitcher, followed by CLAHE filtering and 3D volume stitching in SOFIMA to generate a single stitched montage tomogram. Cryo-CARE denoised stitched montage tomogram is rendered using solved meshes from CLAHE-filtered stitched montage tomogram. Semi-automated actin filament tracing is performed using cylinder-correlation and trace-correlation-line workflow in Amira. After removal of false positives and the filament coordinates are exported for downstream quantitative analysis. The workflow is illustrated for cell2 monatge#1 and the 1.350nm slice through cryo-CARE individual and stitched montage tomogram are shown. Scale bars, 100 nm for individual tomogram tiles and 2 µm for the stitched montage tomogram and filament-trace overview.

